# Exosome secretion kinetics are controlled by temperature

**DOI:** 10.1101/2022.07.22.501177

**Authors:** Anarkali Mahmood, Alan W. Weisgerber, Zdeněk Otruba, Max D. Palay, Melodie T. Nguyen, Broderick L. Bills, Michelle K. Knowles

## Abstract

When multivesicular endosomes (MVEs) fuse with the plasma membrane, exosomes are released into the extracellular space where they can affect other cells. Whether exosomes regulate cells nearby or further away depends on whether they remain attached to the secreting cell membrane. The regulation and kinetics of exosome secretion are not well characterized, but probes for directly imaging single MVE fusion events have allowed for visualization of the fusion and release process. In particular, the design of an exosome marker with a pH sensitive dye in the middle of the tetraspanin protein CD63 has facilitated studies of individual MVE fusion events. Using TIRF microscopy, single MVE fusion events were measured in A549 cells held at 23-37°C and events were identified using an automated detection algorithm. Stable docking precedes fusion almost all of the time and a decrease in temperature was accompanied by decrease in the rate of content loss and a decrease in the frequency of fusion events. The loss of CD63-pHluorin fluorescence was measured at fusion sites and fit with a single or double exponential decay, with approximately 50% of the events requiring two components and a plateau because the loss of fluorescence was typically incomplete. To interpret the kinetics, fusion events were simulated as a point source release of tethered/untethered exosomes coupled with the membrane diffusion of CD63. The experimentally observed decay required three components in the simulation: 1) free exosomes, 2) CD63 membrane diffusion from the endosomal membrane into the plasma membrane at a rate of 0.038 µm^2^/s, as measured by FRAP, and 3) tethered exosomes. The final component of the decay arises from exosomes being secreted but tethered to the surface with one tether that has a lifetime of 8 seconds at 37°C and longer at lower temperatures. Simulating with fixed tethers or the absence of tethers fails to replicate the experimental data. This kinetic analysis increases our understanding of exosome secretion and how it is regulated by temperature. Our model suggests that exosome release from the fusion site is incomplete due to post-fusion, membrane attachment.

## Introduction

Exosomes are a subset of small extracellular vesicles (sEVs) secreted from cells. They range from 30-100 nm in diameter and are formed from the inward budding of vesicles into the intraluminal space of late endosomes^1–3^. Exosomes are secreted into the extracellular space when these multivesicular endosomes (MVEs) fuse with the plasma membrane^1,4–6^. Once exosomes are secreted into the extracellular space, they can affect cells nearby and further away^7–9^. Exosome secretion is used by healthy cells to maintain essential processes such as homeostasis^10^ and cell motility^11,12^, however the release of exosomes can be exploited by unhealthy cells to aid in disease progression^13,14^. Exosomes likely facilitate disease progression via the transfer of biomolecules from unhealthy to healthy cells^1,15–18^. Specifically, exosomes and other sEVs have been shown to play a role in neurodegenerative diseases^19–21^ and cancer^7^. Many studies regarding disease states isolate sEVs, which are rich in exosomes, but not exclusively exosomes therefore the term “sEV” is used. The discovery of exosomes and their involvement in disease states has motivated research into exosomes as markers for early detection and potential avenues for intervention.

Though MVE membrane fusion is likely an integral check point that can be harnessed to modulate exosome release, a mechanistic understanding of the fusion process is lacking. It is known that MVE fusion is a constitutive process but enhanced in the presence of Ca^2+ 16,22^. In a bulk sEV collection assay, more sEVs were collected from cells treated with Ca^2+^ ionophores^16^. This was also observed in single fusion events, where ionophores increased the number of events^22^, however the magnitude of the effect of Ca^2+^ depended dramatically on the cell type^22^. Basic information, that can further our understanding of the fusion process, including whether MVEs are stably docked, identification of proteins that control fusion and the dependence of MVE fusion on temperature are unknown. This information can provide insights into similarities and differences between well-understood fusion mechanisms utilized by other vesicles, such as dense core vesicles, and potentially identify avenues for modulating exosome secretion.

Membrane fusion has been well-studied regarding the regulated fusion of secretory vesicles, such as synaptic vesicles, dense core vesicles, and insulin granules. By analyzing the kinetics of content release during fusion between secretory vesicles and the plasma membrane has led to the identification of different fusion mechanisms, such as kiss and run^23^ and a diversity of fusion modes that affect the release of content^24^. As kinetic studies of different vesicles have provided highly important insights into stimulated exocytosis, similar studies regarding MVEs fusion will be essential to increase our understanding of exosome release.

To elucidate the kinetics of exosome secretion, a marker of intraluminal vesicles (ILVs), CD63, has been used to visualize single fusion events^25–27^. CD63 is a tetraspanin protein present on the endosome and plasma membranes of cells and enriched on the membrane of ILVs that become exosomes upon MVE fusion^16^. Several labs have tagged CD63 on the first extracellular loop with a pH-sensitive fluorescent protein (*i*.*e*. pHluorin or pHuji) such that the probe is quenched on the inside of the late endosome or when on the surface of an ILV^25–27^. CD63-pHluorin is a pH dependent probe, that remains quenched inside the acidic environment of an MVE and signals the onset of fusion by a sudden spike of fluorescence that gradually disperses radially. The rate at which fluorescence dissipates depends on how CD63 labeled exosomes leave the fusion site. Post fusion, exosomes have been observed to either diffuse far from the site of secretion^9^ or remain close to the cell surface^28,29^. Interestingly long lasting fluorescence after fusion has been observed with CD63+ and CD81+ exosomes but not CD9+ exosomes^27^. Tethering molecules, such as tetherin, may be involved with limiting the widespread release of a portion of exosomes^30^. By observing the kinetics of single fusion events, we demonstrate that the release of exosomes from the fusion site is incomplete and simulations of the process allow us to propose that this is due to the extent of exosome attachment to the cell surface.

In this work, single MVE fusion events were visualized using CD63 fluorescent probes and total internal reflection fluorescence (TIRF) microscopy. A non-small cell lung cancer cell line (A549) was used as a model system to investigate MVE fusion because they readily release exosomes, facilitating the imaging process, and exosomes have been shown to play an integral role in cancer progression^14,31–33^. One challenge of measuring constitutive fusion events in a single vesicle fusion assays is the tedious process of manually finding and analyzing fusion events that occur at random points in time and in a relatively slow fashion (∼1-3 events per minute). To overcome this, an automated approach to detection and analysis capable of capturing both large and small fusion events that occur on top of a background of CD63-pHluorin on the plasma membrane was developed. Our results reveal that, prior to fusion, almost all MVEs are docked for at least 1 second, typically longer. The frequency of fusion increases and the kinetics of release is faster at higher temperatures. The kinetics of release, or loss of fluorescence post-fusion, can relay information about the fate of exosomes. Through two types of analyses, 1) fitting the fluorescence loss and 2) simulations of the release event, we determine that a portion of exosomes are free to diffuse away from the fusion site, but some exosomes remain attached to the surface in many, but not all, fusion events. This attachment depends on temperature, with fewer exosome remaining attached at higher temperatures.

## Materials and Methods

### Cell Culture

A549 cells were cultured in Dulbecco-modified Eagle’s minimum essential medium (DMEM; Gibco 11965092). DMEM was supplemented with 10% fetal bovine serum (Sigma Aldrich). Cells were grown and maintained in a humidified 37°C, 5% CO_2_ incubator. Cells used for microscopy were plated in LabTek 8 well dishes and were transiently transfected using Lipofectamine 3000 (ThermoFisher) using 2.5 mg/ml of CD63-pHluorin plasmid DNA (gift from D.M Pegtel^27^) according to the manufacturer’s protocol. Cells were imaged between 24-48 hours post transfection.

### Total Internal Reflection Fluorescence (TIRF) Microscopy

The transiently transfected cells were imaged using an inverted Nikon microscope equipped with 491 nm and 561 nm lasers on an acousto-optic laser launch (Solamere Technology) and an EMCCD camera (Andor iXon897). The laser power entering the back of the microscope was set to 30 mW for each laser and then reduced using ND filters in the laser path prior to excitation. To direct excitation light to the sample, a dual color, a TIRF-specific, dichroic beam-splitter was used (Chroma Technologies) and emission was split into red and green fluorescent channels using a Dual-view (Optical Insights). For pHluorin detection, a green laser (491 nm) was used for excitation and a 525/50 filter (Chroma Technologies) was used to detect emission. For pHuji detection, a yellow laser (561 nm) was used for excitation and a 605/75 nm filter (605/75) was for detection. For magnification, a 60x 1.49 NA objective and an additional 2.5x lens were used such that one pixel is 109 nm. To improve our ability to focus on both colors simultaneously a gtfflong focal length (1000 mm) plano-convex lens was placed in the red emission channel of the dual view. Micromanager image acquisition software^34^ was used to obtain data with a 50-100ms exposure time continuously for 500-1100 frames. Prior to recording cells for two color imaging, the TIRF field was aligned with 200 nm diameter, carboxylate modified, yellow-green Fluospheres (ThermoFisher). Yellow-green Fluospheres show up in both channels and are used for registration of green and red images in conjunction with a custom MATLAB code. The depth of field is also indirectly measured by the Fluospheres and the sample temperature was maintained using an on-stage heater system (Bioscience Tools, TC-1-100S).

### Image Analysis

To identify fusion locations, image sequences with the pH sensitive MVE fusion marker were subject to the following analysis using scripts designed in MATLAB (Fig. S1). The analysis code is available at GitHub (https://github.com/michelleknowles/membrane-fusion)

#### Step 1 - Calculation of Differential Movies

Images were subtracted from one another to highlight the cellular locations where intensity has changed in time. An earlier time frame (t) was subtracted from a later time frame (t + 25 frames, t + 1.25 sec). The maximum projection of the difference movie was used for the subsequent spot finding step. Bandpass filtering of the movie helped with finding spots (Fig. S2) but was not essential and not applied for the fusion events processed here.

#### Step 2 – Fusion Event Localization

The average intensity of the maximum projection is used as the threshold to locate spots and a cell mask is applied to find the average cell intensity. Spots above the calculated threshold were located using MATLAB tracking algorithm originally designed for use in IDL^35^.

#### Step 3 – Cropping raw data at fusion locations

A 25 × 25 pixel region with the potential fusion spot centered was cropped. If two color channels were measured, both were cropped.

#### Step 4 – Output the intensity in time

The average intensity from a circle 7 pixels in diameter was measured. The average cell intensity within the cell boundary was defined as the background for fusion events and subtracted from fusion event. Intensity traces were normalized by first subtracting the minimum before fusion then dividing by the maximum intensity.

#### Step 5 – Alignment of data in time

Since the fusion events are not synchronous, all events were aligned by setting the frame prior to fusion to be 0 s. A background average is calculated using first 35 data points of the fusion event intensity data and the onset of fusion is defined as the point where intensity spikes to 1.4 times the background average.

#### Step 6 – Sort events

Events that are identified in the above steps are either fusing, moving or docking vesicles. Initially, events were sorted by viewing the cropped movies prepared in Step 3 or by the intensity traces in Step 5. Fusion is noted by a sudden spike in intensity, which is followed by gradual loss of fluorescence over time. With a subset of vesicles, an automated protocol was developed to separate fusion from other events. Here, docking events were removed based on the slope of the decay after the maximum intensity. Tracking was performed on a subset of the potential fusion events and the rate of motion was used to filter out fusing vesicles from moving vesicles. Moving vesicles are removed based on the mean square displacement and a diffusion coefficient greater than 0.010 µm^2^/s denoted a motile vesicle. These non-fusion events are shown in Fig. 2.

#### Step 7a – Calculate kinetic parameters

From the aligned intensity traces in time several pieces of information are noted in the results. The slope of the decay, fraction lost at 1 and 5 seconds was calculated. To determine the rate of decay, a double exponential decay shown in Eq. 1, was used to fit single events from the maximum intensity to the end of the trace.

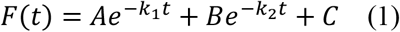

Where A and B are the relative amounts of the fast and slow components that have corresponding rates k_1_ and k_2_. C is a plateau because some traces do not return to the background level over the time intensity was measured. The rates from fitting, k, were converted in t_1/2_ where t_1/2_= ln(2)/k. All fitting was performed in GraphPad Prism.

#### Step 7b – Calculate size changes in time

The radial plot of the fusion event was determined as a function of time; fusion events expand in time^36^. To determine the size of the fusion events, the cropped 25×25 pixel image at the moment of fusion (t = 0) was averaged with the frame before and the frame after (t = −0.05 to 0.05s). This was done to improve the signal to noise. A radial plot was calculated from the image such that the center pixel is position 0 and the equidistant pixels are averaged, as described in our past work^37^. The FWHM was determined by linear interpolation between the two points surrounding the half maximum intensity.

#### Two-color TIRF imaging of docking and fusion

To determine whether MVE vesicles dock prior to fusion, A549 cells were co-transfected with EGFP-CD63 and CD63-pHuji. CD63-pEGFP C2 was a gift from Paul Luzio (http://n2t.net/addgene:62964; RRID:Addgene_62964) and CD63-pHuji was a gift from D.M. Pegtel^27^. Two color TIRF microscopy was performed followed by automated fusion detection using the pHuji channel. To count whether the MVE was docked prior to fusion, the intensity before fusion in the green channel was measured for events where green fluorescence was observed during fusion, which ensured that CD63-EGFP was present in the MVE as some events could be noted as not docked prior to fusion due to the absence of CD63-EGFP in that particular MVE. Presence of EGFP-CD63 at the fusion site was confirmed both manually using EGFP-CD63 images and via the small increase in EGFP-CD63 brightness.

### Fluorescence Recovery After Photobleaching (FRAP)

FRAP was performed on A549 cells expressing CD63-pHluorin using an Olympus Fluoview 3000. Cells expressing CD63 on the plasma membrane were selected and a 1.99 µm radius spot was bleached using the tornado raster setting. Three images were collected prior to bleaching and images were collected for a total of 100 frames by taking one frame every 1.085s. The rate of recovery was fit, as described previously, to determine the diffusion coefficient of CD63 on the plasma membrane and the fraction mobile^38^. FRAP was performed at 23°C and 37°C.

To measure the time, t_1/2_, it would take for half the particles moving at the rate measured in FRAP to leave a 0.76 µm diameter circle we calculated the probability distribution of step sizes taken in space and time^39^,

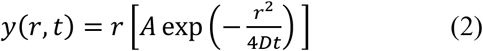

and solved for t_1/2_, the time where half of the molecules are within a circle of radius, r = 0.38 µm, and half have traveled further. D is the diffusion coefficient (0.039 µm^2^/s) of CD63 on the plasma membrane yielding t_1/2_ = 1.33s. This was verified by simulations of a random walk, described below.

### Modeling of MVE Fusion Decays

MVE fusion events were modeled as a point source where 100 CD63-pHluorin molecules are deposited. Post deposition, molecules can escape a circle of 0.76 µm diameter (7 pixels in the image analysis) around the fusion site in several ways: 1) As an untethered exosome moving at 6.5 µm^2^/s. This rate of diffusion is calculated using the Stokes-Einstein diffusion equation: D = k_B_T/(6πηr), where k_B_ is the Boltzmann constant, T is 310 K, η is the viscosity of the aqueous buffer (0.69 cP), r is the radius of an exosome, which ranges from 15-60 nm and 50 nm was used. The rate an exosome would diffuse from the site of fusion varies by less than 10% over the range of temperatures measured in the experimental data. 2) As a molecule diffusing from the endosomal membrane into the plasma membrane at 0.039 µm^2^/s. This rate of diffusion comes from the FRAP measurements of CD63-pHluorin diffusing on the plasma membrane of live A549 cells. 3) As tethered exosomes. The rate of motion of the tethered exosome is immobile unless the tether is broken, then the exosome diffuses at a rate of 6.5 µm^2^/s. In the simulation, the time constant at which half of the tethers are broken (t_1/2tether_) was varied from 1-50 seconds to best fit the data and exosomes were attached with only one tether. The fraction of CD63-pHluorin in the membrane, in tethered exosomes, and in free exosomes was a second variable that was altered to best match the data. The simulation used 50 µs steps and allowed the molecules to move via diffusion with the rates above if untethered and checked at each 50 µs time point if the tether was still intact. The location of the molecule was recorded every 50 or 100 ms, depending on the experimental data that the simulation was matching to; 23°C data was taken at 100 ms/frame to obtain longer data sets due to the observed plateau in the events and the rest was taken at 50 ms/frame. The simulation lasted between 12.5 and 25 seconds and the number of molecules remaining in a 0.76 µm circle, the size over which the intensity was measured for the experimental data analysis, was output.

Simulations were run and a best match to the average decays was obtained for all temperatures and individual decays for 37°C data (n = 98 fusion events). For the average data, the ratio of free exosomes to endosomal CD63 diffusing on the membrane was determined by fitting the first 10 frames of the experimental data. The tethered exosomes do not contribute to intensity loss at this stage. Next, the t_1/2tether_ was varied and the percent of tethered exosomes was determined to fit the long-time tail of the experimental data. At this point, the endosomal CD63 and the free exosomes do not contribute significantly to the experimental data. For fitting 5 simulations were performed and averaged. For individual traces, the initial 10 frames were fit as described for the average data. The t_1/2tether_ was kept constant after being determined from fitting the average, but the fraction tethered was varied in 2% increments from 0 to 100%. The remainder of the CD63 was split between endosomal and free exosomes at a ratio of 0.6, as determined from fitting the averages. The absolute value of the differences between the simulation and the data was measured for each time point and the lowest sum of these differences was considered the best match for both average and individual traces.

## RESULTS

### Automated detection and quantification of MVE fusion events

MVE fusion events were visualized using TIRF microscopy and quantified in A549 cells using CD63-pHluorin, a pH-sensitive fluorescent probe^27,40^, where the pHluorin moiety is located between two transmembrane domains of CD63 such that it is localized to the interior of the MVE and on the exterior of the ILV (Figure 1A). While enclosed in the acidic environment of an MVE, pHluorin is quenched. Once the vesicle fuses with the plasma membrane, the pH change unquenches pHluorin leading to an abrupt fluorescence spike, marking the onset of MVE fusion (Fig 1B-D). The fluorescence rapidly appears at fusion site and dissipates into the surrounding area (Fig 1D), as exosomes diffuse into the extracellular space and as CD63-pHluorin on the MVE membrane diffuses into the plasma membrane. The kinetics of MVE fusion can be measured from the intensity profile of single events in time (Fig 1C).

**Figure 1:**
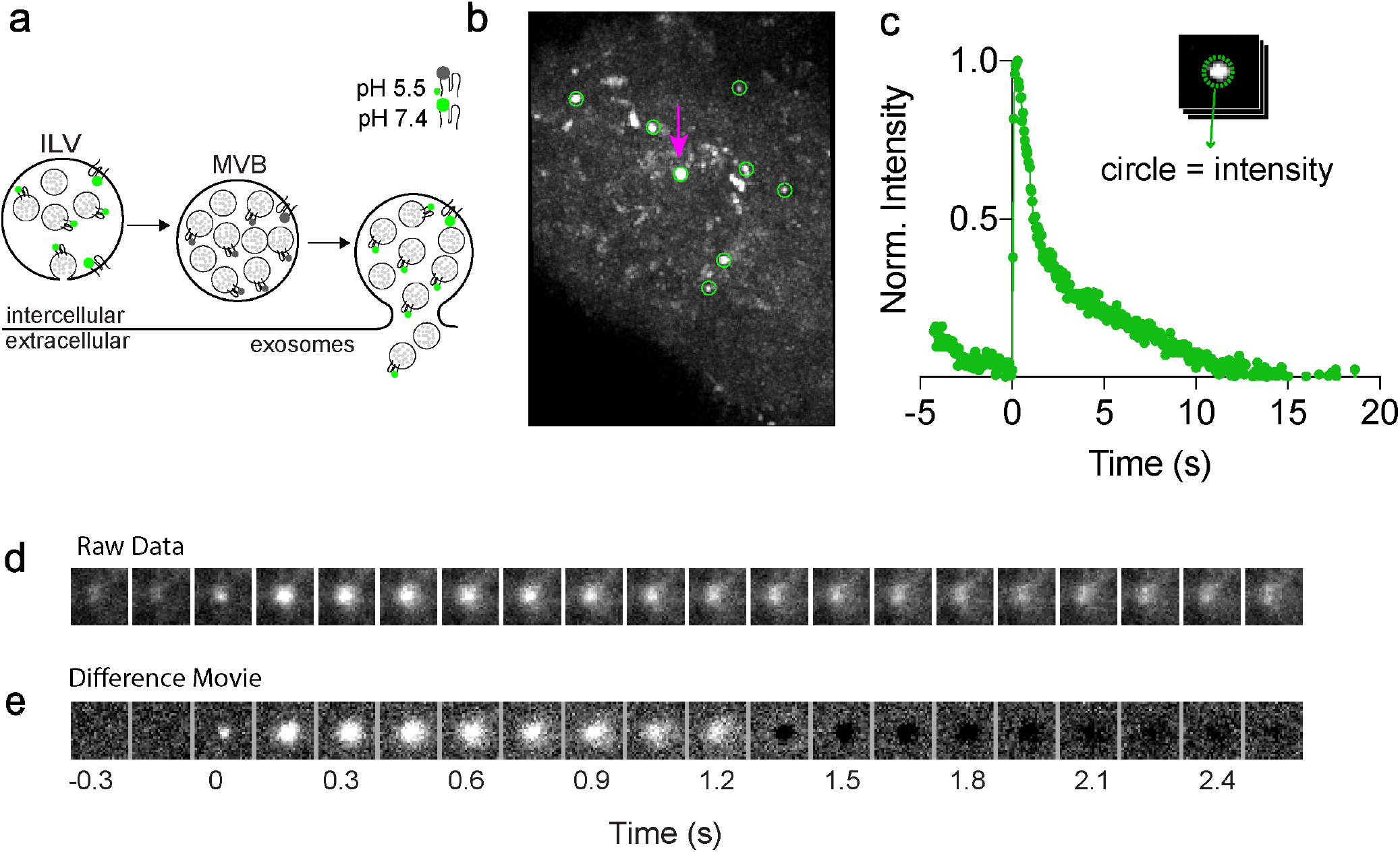
MVE fusion events are visualized using CD63-pHluorin. a) Diagram depicting the assay used to detect MVE fusion events. Cells are fluorescently labeled with CD63-pHluorin, which is quenched when in an acidic vesicle. Once the MVE fuses, the pH increases and pHluorin emits green fluorescence. b) Membrane fusion is observable in live cells using TIRFM. A maximum projection of a band pass filtered, difference movie of an A549 cell with potential fusion events encircled in green, scale bar = 5 µm. A total of 8 events were found, shown in green circles c) Example of a single exocytosis event, where 0 s is defined as the onset of fusion. Scale bar = 1 µm d) The intensity profile of a sample fusion event showing a rapid increase in fluorescence upon MVE fusion. The intensity shown is the average intensity of one fusion event within a 0.76 µm diameter circle and normalized to the maximum intensity. d) Single images of raw data for a fusion event. e) Single images of a difference movie. Maximum projections of difference movies were used to identify fusion locations. (Cell AM1587)

Although sites of fusion are visible by eye, manually locating and detecting each event is time consuming and subjective. To circumvent these issues, event detection was automated in MATLAB and the procedure is summarized in Supplementary Fig S1. Detection begins by enhancing the signal of the fusion events relative to the background via the generation of a difference movie, like others have done for constitutive fusion measurements^41^. The purpose of this step is to remove signal rendered by vesicles that are visible and stationary on the cell surface throughout the movie; fusing vesicles show a rapid fluorescence intensity spike due to pH change upon fusion and are generally not visible before the onset of fusion (Fig 1D). The difference movie is calculated by subtracting an earlier frame from a later frame. Examples of the effects of different time intervals and filtering on the signal to noise of fusion events are shown in Supplementary Fig S2. Signal enhancement obtained from the difference movie is apparent in Fig 1E, a single fusion event that has been bandpass filtered, and Supplementary Fig S2, which shows an example of a single image (Fig S2A), the maximum projection before (Fig S2C) and after (Fig S2B) the difference movie calculation. When a difference movie is not used, many more potential fusion locations are identified (Fig S2C) however, these are primarily stationary vesicles present throughout the movie. Potential fusion events are then detected using a maximum projection image of the difference movie (Fig S2D). Once located, potential fusion sites were cropped from the raw movie file into 25 × 25 pixel regions centered around the fusion spot and the fluorescence intensity within a 7 pixel (0.76 µm) circle, centered on the fusion spot, was quantified for each fusion event and output as a function of time (Fig 1C). Fluorescence intensity traces in time are used to identify fusion events and then further analyzed for kinetic information.

Upon the fusion of an acidic vesicle, the fluorescence intensity profile of a pH dependent probe exhibits certain criteria, such as a rapid increase in fluorescence intensity at a localized spot followed by a cloud-like spread of intensity into a region around the initial spike location (Fig 1D)^27,36,41,42^. Therefore, fusing MVEs were identified by the rapid onset (1-2 frames) of fluorescence intensity, followed by a gradual loss of fluorescence over time. Our approach to locating the fusion events works well for avoiding stationary, fluorescent vesicles, however, fluorescent vesicles that move or dock are detected. Figure 2 shows the most common examples of non-fusion events detected. Among the potential fusion events identified using the automated approach, approximately 70% were fusion events (Fig 2A). The remaining 30% of non-fusion events comprised of a combination of moving vesicles (Fig 2B) and docking vesicles (Fig 2C). While all three events show a rapid onset of fluorescence in the intensity trace, the subsequent kinetics are different. Fusion events decay exponentially (Fig 2A). Moving vesicles typically have to remain stationary for a few frames to be detected as a spot, then move away causing a slower onset of fluorescence followed by a plateau or slow decay (Fig 2B). Docking events do not decay, resulting in a fluorescence plateau (Fig 2C). The slope from the maximum intensity to one second later was fit and used to bin the events into fusing, moving, and docking (Fig 2D). While slope is useful for binning docked vesicles from the others, significant overlap was observed for fusing and moving. Therefore, tracking was performed to determine a diffusion coefficient (Fig 2E); moving vesicles can be identified by a diffusion coefficient greater than 0.010 µm^2^/s. Once the fusion events were isolated from other detected events, the intensity traces were further analyzed to characterize MVE size and MVE fusion kinetics.

**Figure 2:**
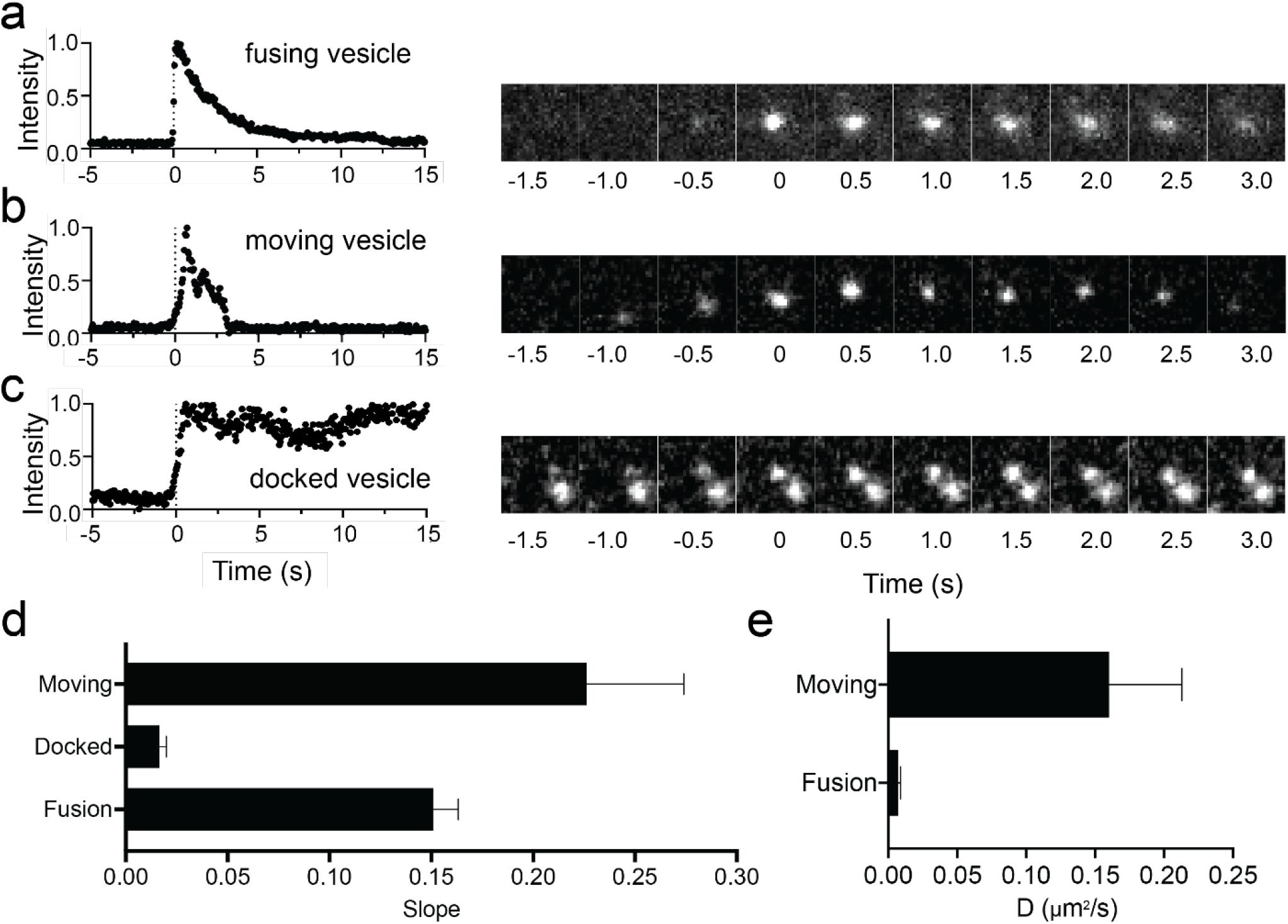
Potential fusion events can be categorized as A) vesicle fusion, B) vesicle motion or C) vesicle docking. D) Docking is noted by the lack of loss of fluorescence in the intensity trace and slow onset. The slope of a line fit from the maximum to one second later is substantially lower (N = 15 events each), E) The moving vesicles can be separated based on diffusion coefficient from tracking analyses (N = 15 vesicles each). Fusion events are then binned and separated for analysis.

Following the fusion event onset, CD63-pHluorin gradually moves away from the fusion site and a radial spread of the fluorescence is typically observed (Fig 1D and 2A). This presumably occurs as exosomes diffuse from the fusion site and the CD63-pHluorin from MVE surface laterally diffuses into the plasma membrane as depicted in Fig 3A. Using the images of a fusion event (Fig 3B), radial averages were calculated from the site of fusion (the center of the image), in time (Fig 3C) and compared to the size of docked, dense core vesicles in PC12 cells and 200 nm diameter yellow-green fluospheres (Fig 3D). Quantification of the width of the radial plots at the onset of fusion (t = 0 s) yielded an average vesicle diameter of 440 ± 11 nm (Fig 3E) and an increase in size, denoted by the black dashed lines compared to the size at the start (blue dashed line), was observed as the event progressed (Fig 3C). The width of the intensity at the fusion site increased in time due to the loss of CD63-pHluorin from the fusion site and into the surrounding area.

**Figure 3.**
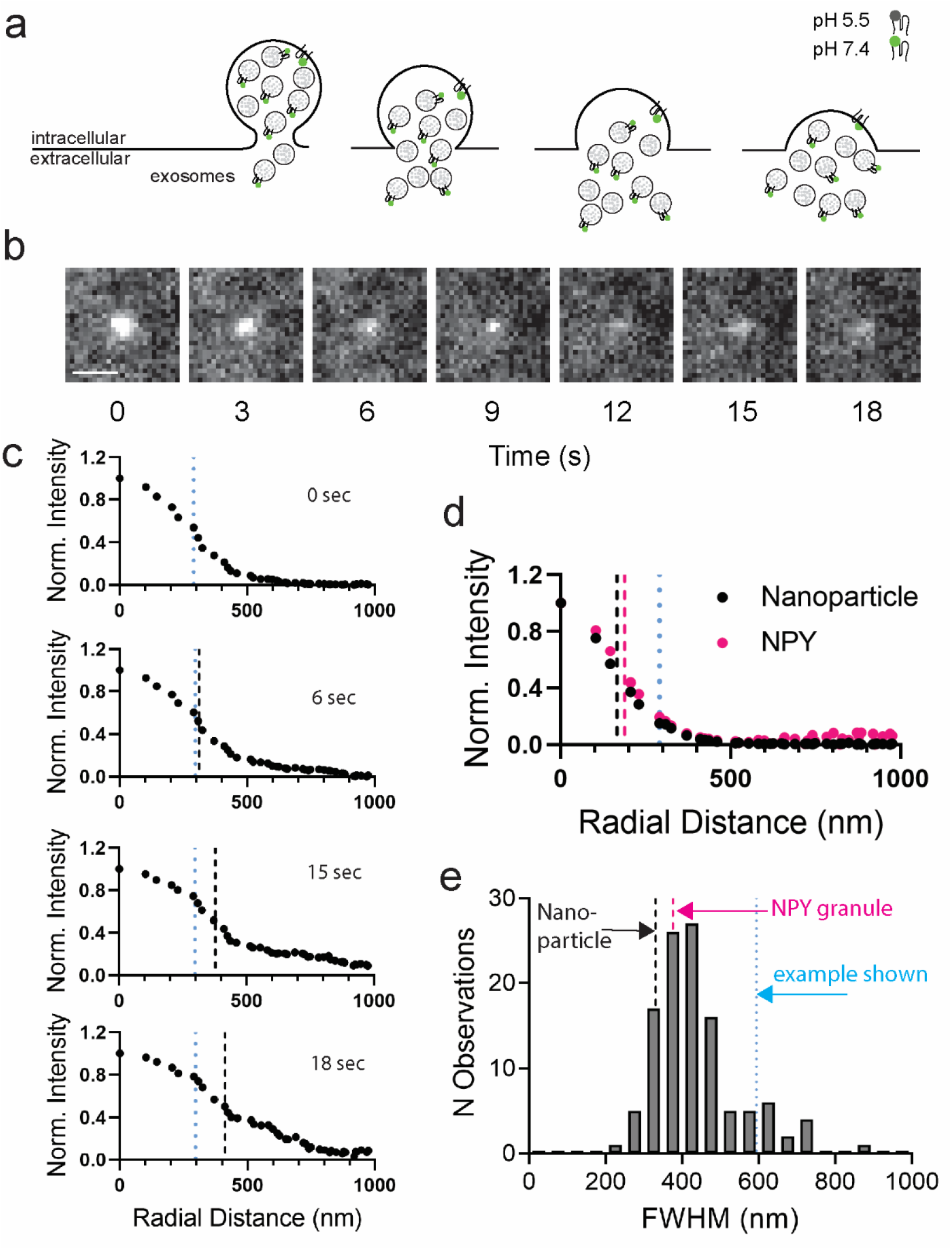
MVE fusion events expand in time. A) Depiction of MVE fusion showing how CD63+ exosomes and CD63 in the membrane lead to spread of the fluorescence signal in time. B) Sample fusion event, unfiltered with contrast the same for all images. Scale bar is 1 µm. C) Average radial plots of the fusion event showing a change in MVE radius (HWHM) over time. The blue dashed line is the initial radius (t = 0s) and the black dashed line is the radius at that point in time. D) For comparison, the size of NPY-mCherry granules in PC12 cells and yellow-green fluorescent nanoparticles (d = 200nm). The blue dashed line is the HWHM shown in C. E) Histogram showing range of MVE diameters (FWHM) at the onset of fusion (t = 0 s).

### MVE fusion leads to exosome release and deposition of CD63 on plasma membrane

By observing individual fusion events, the rate of the loss of intensity in time was analyzed to determine the mechanism by which CD63-pHluorin leaves the fusion site. Heterogeneity in the rate of fluorescence decay was noted with two types of decay profiles (Fig 4A-B); the decay profiles of some secretion events underwent a biphasic decay with an initial fast downward slope for approximately 3s, followed by a gradual intensity loss phase (Fig 4A), whereas others showed a gradual loss of intensity signal (Fig 4B). The average of 110 events is shown in Fig 4C and many single fusion events taken from two cells are shown in Supplemental Fig 3A. A majority of the fusion events exhibited two-component decay profiles (Fig 4D) at 37°C, however, a few events were not able to be fit well if the movie ended before a long-time decay could be observed (see Fig S3 for examples).

**Figure 4.**
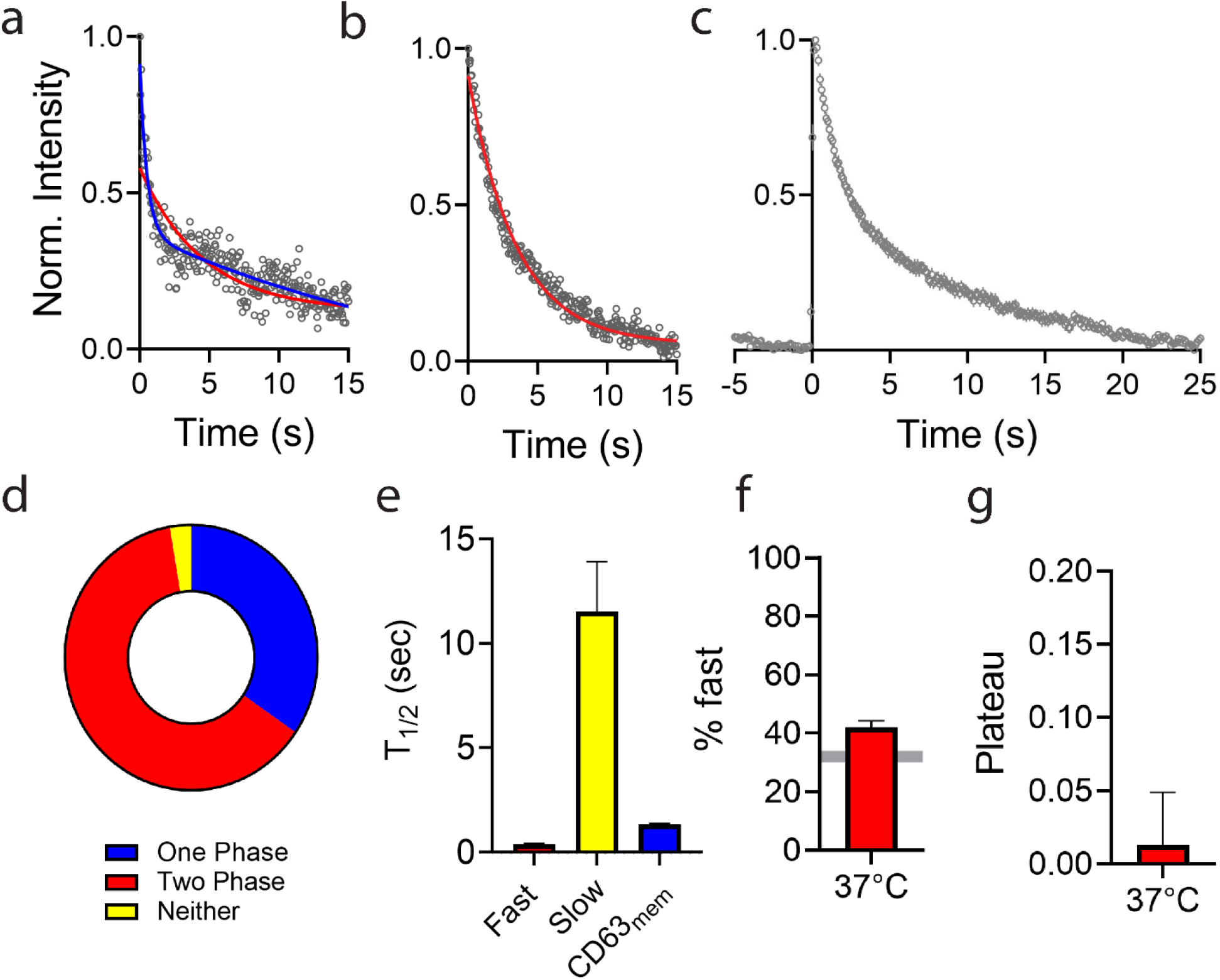
Multiple kinetic modes are observed in MVE fusion events. A) A single fusion event (black dots) fit to a one component (red) or a two component (blue) fit. B) Sample fusion event that is fit well with a one component decay. C) Average time course of 120 fusion events. D) Pie chart showing percent of one vs two component decay profiles. E) Average fast (red) and slow (yellow) decay constants where t_1/2_ = ln(2)/k, calculated using the rate constants from bi-exponential decay fits of individual fusion events. The mobility of CD63-pHluorin on the plasma membrane was measured by FRAP and an expected t_1/2_ was calculated based on the diffusion coefficient and circle size (see Methods). All data taken at 37°C. F) Percent of the fast component in decay curves for fusion events at 37°C (Average +/− SEM). G) Plateau for decay curves at 37°C (Median +/− 95% CI).

To determine the cause of the biphasic decays, we considered the likely locations of the fluorescence probe. CD63-pHluorin, is present on the ILV and the MVE surface. Recent EM data shows that, on average, 30-34% of CD63 is retained on the endosomal limiting membrane, whereas the remainder is located on ILVs^43^. Indeed, CD63 was present on collected exosomes (Figure S4). When fusion occurs, it is expected that part of the decay is due to exosomes diffusing from the fusion site and the part of the decay is due to the CD63-pHluorin diffusing from the MVE membrane into the plasma membrane. Analysis of the rate of decay was performed to identify the amount of fluorescence loss due to each of these components. The traces that fit well to a two-component exponential function (n = 66 events), displayed fast and slow components with t_1/2_ of 0.40 ± 0.04 s and 11.5 ± 2.4 s, respectively. To determine which component of the kinetics was due to CD63-pHluorin diffusion in the plasma membrane, fluorescence recovery after photobleaching (FRAP) experiments were performed with CD63-pHluorin on the surface of A549 cells (Supplementary Fig S5). Recovery traces (Fig S5B) show that CD63-pHluorin is mobile on the plasma membrane and fitting of the data revealed that CD63-pHluorin diffused at a rate of 0.039 µm^2^/s (Fig S5D). Interestingly, there was no temperature dependence observed for the motion of CD63 on the plasma membrane (Fig S5). To determine how long it takes for molecules to leave a region the size (*d* = 0.76 µm) over which intensity was measured for a fusion event, data was simulated for a 2D diffusion of particles moving at 0.039 µm^2^/s and deposited at the center of the circle. On average, half of the particles are lost from the fusion site at 1.3 +/− 0.1 s (Fig 5E, blue). The fraction of the intensity that moves at the faster rate is 40%, slightly higher than the expected amount to be present on the endosomal membrane, 30-34%^43^, denoted by the grey bar. The plateau, although small, was above 0 (Fig 4G) as fluorescence often remained post fusion.

**Figure 5:**
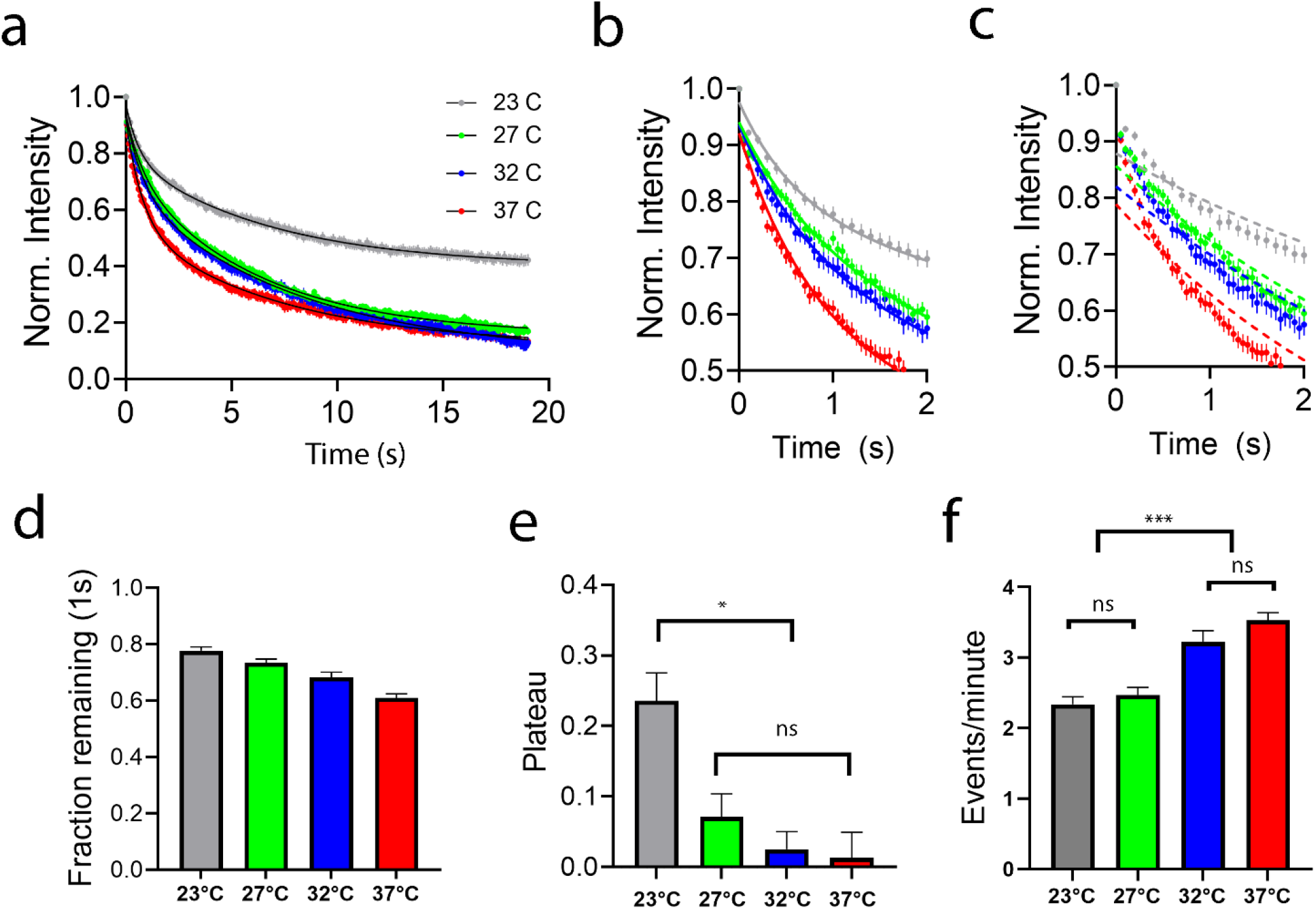
Kinetics of single MVE fusion events depend on temperature. a) Average intensity traces in time of single MVE fusion events at 23°C (n=86), 27°C (n=83), 32°C (n=77), and 37°C (n=110). For fitting, the time alignment was done with respect to the maximum intensity (0s) and individual traces were normalized by the maximum. Solid lines are fits with a biexponential function, b) zoom in on the biphasic fit and the c) single exponential fit at short times. Error bars are SEM. d) The percent loss in intensity one second after the maximum. All are significantly different in t-tests (p > 0.05) from the nearest temperature (Average +/− SEM). e) The plateau from the biexponential fit relates the long time, remaining intensity (median +/− 95%CI, only 23°C is significantly different from others). f) Fusion events observed per minute of data acquisition at different temperatures (Average +/− SEM).

### MVE fusion is temperature dependent

MVE fusion kinetics were analyzed at four different temperatures: 23°C, 27°C, 32°C, and 37°C. The kinetics of MVE fusion are directly related to temperature; a decrease in temperature caused a decrease in the rate at which the CD63-pHluorin fluorescence signal was lost (Fig 5A-D). Although the lower temperatures reduced the rate of content loss, two phases were required when fitting the average decay. In Fig 5B, the decays are fit well with a bi-exponential but the single exponential fit in Fig 5C clearly misses the fast component for all temperatures. This suggests that the release event contains at least two mechanisms of leaving, likely the loss of CD63 on the endosomal membrane and loss of exosomes themselves, occur for all temperatures studied. The fast portion of the decay lasted about 0.4 s for 37°C (Fig 4E) therefore, we compared the amount of fluorescence remaining after 1s for all of the different temperatures to isolate the fast component. The decays after 1 second trend as a function of temperature (Fig 5D), however the mobility of CD63 on the membrane shows no temperature dependence in FRAP measurements (Fig S5), suggesting that the exosome release is the temperature dependent portion, not the diffusion of CD63 through the membrane.

Reduced temperatures also led to the incomplete loss of fluorescence from the fusion site. The remaining fluorescence in the plateau was much more prominent at lower temperatures (Fig 5E) with the average of the observed events at 23°C retained approximately 50% of the maximum intensity at 10 s post fusion, whereas the average of the fusion events observed at 37°C only retained 50% of maximum intensity for approximately 1-2s post fusion (Fig 5A). The plateau observed at 23°C was approximately 15 times larger than the plateaus observed at 23°C and 3-5 times larger than 27°C or 32°C. The presence of a plateau suggests that content is not fully released from the fusion site or remains attached to the cell surface and higher temperatures allow more content to leave the fusion site. The rate of fusion events per minute (Fig 5F) also trended higher at higher temperatures. Overall, higher temperatures make fusion more frequent, faster to release content and less fluorescence is retained at long times.

### Simulation of exosome release, tethering, and CD63 membrane diffusion can account for the experimentally observed kinetics

Fitting the data to an exponential function has limitations, due to the signal to noise and total duration of recordings available when measuring single fusion events (Supplementary Fig S3 for examples). Therefore, we applied a simple, physical model to determine what types of CD63 motion were sufficient for explaining the release kinetics at different temperatures. In the experimental data, the loss of signal at the fusion site is due to CD63-pHluorin leaving. It is likely that CD63-pHluorin can leave in several ways: 1) CD63-pHluorin is sorted onto ILVs that become exosomes and these exosomes can diffuse freely from the fusion site. 2) CD63-pHluorin exosomes can remain at the fusion site due to tethering^30^. 3) CD63-pHluorin is on the endosomal membrane and can diffuse into the plasma membrane to leave the fusion site. To simulate the data, molecules were released at the center of the fusion site, which was defined as the center of a 0.76 µm diameter circle (Fig 6A). The molecules move with one of the three motions above. Free exosomes (Fig 6A-B, green) diffuse away almost instantly at a rate of 6.5 µm^2^/s. CD63-pHluorin located on the endosomal membrane (Fig 6A-B, blue) moves at a rate of 0.039µm^2^/s as measured by FRAP of CD63-pHluorin on the plasma membrane (Fig S5). Finally, tethered exosomes (Fig 7A, red) do not move. To account for the long-time (t > 5s) decay observed in the data (Fig 6B, grey open circles), we considered that the intensity loss could be due to photobleaching of CD63-pHluorin. Photobleaching was measured (Fig 6B, grey open squares) and the rate of photobleaching is extremely slow under the imaging conditions, much slower than the loss of fluorescence from the fusion site at long time (Supplementary Fig 7A-B). Therefore, to account for the long-time decay observed, tethers were allowed to break, and without breakage, the long-time data could not be replicated (Supplementary Fig 7C-D). Therefore, tethers were considered breakable with some time constant, t_1/2tether_. The newly released exosomes are shown in Fig 6A (red with black outline) and allowed to diffuse at the rate of a free exosome when tethers break. In all simulations, the amount of CD63 remaining in the 0.76 µm circle was measured and an average of 5 or more simulations, each with 100 molecules, of these three dynamic components is shown separately in Fig 6B alongside the experimental data at 37°C and the photobleaching loss.

**Figure 6:**
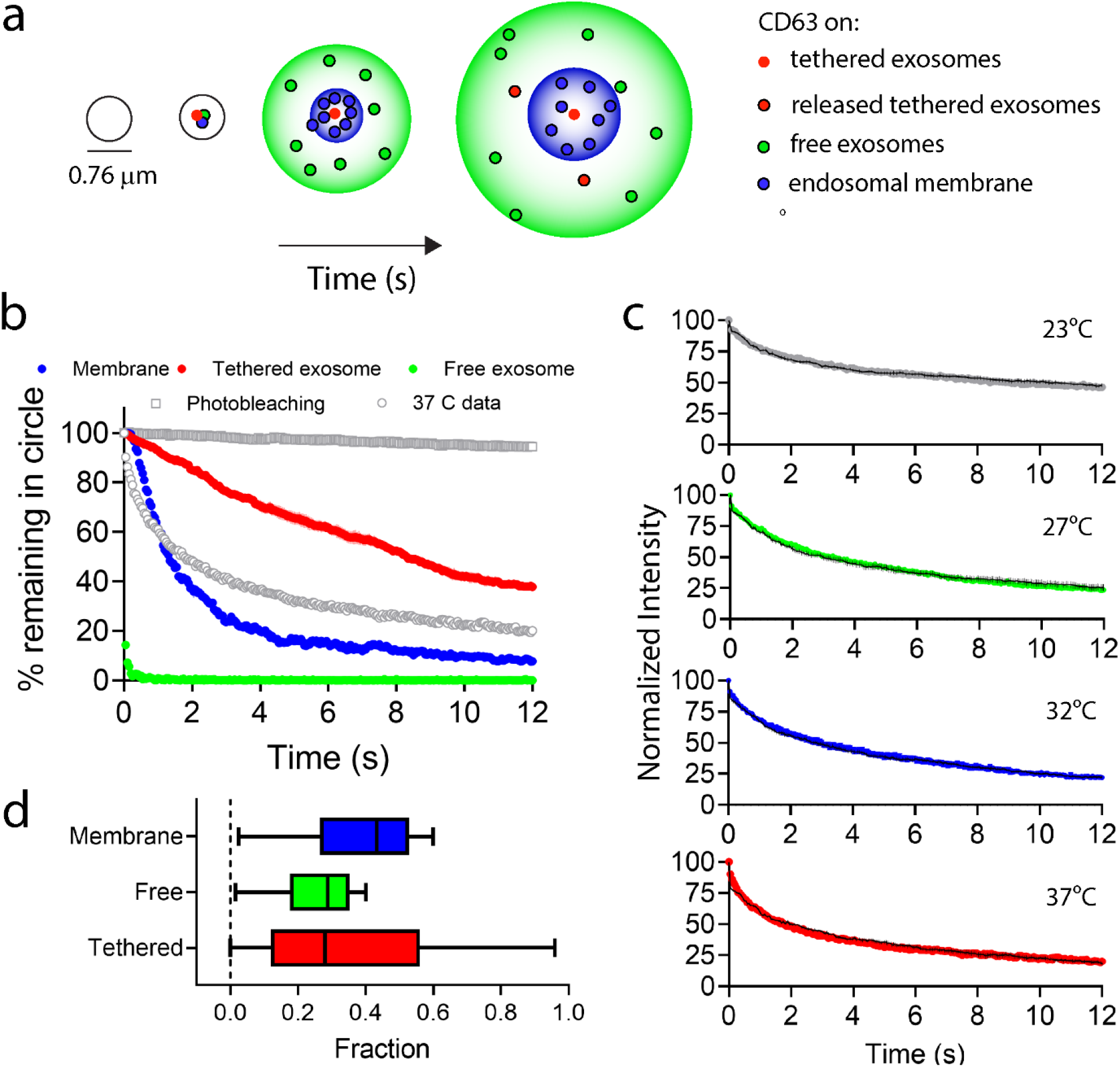
Fusion events were simulated by allowing CD63 particles to leave the center of the fusion site as tethered exosomes, free exosomes or membrane diffusion. a) depicts the simulation in time. 100 particles are deposited at the center of the 0.76 µm circle and allowed to diffuse unless they are tethered. A portion of the particles are freely diffusing as 100 nm diameter exosomes (green), diffusing on the plasma membrane (blue), or tethered, which are not mobile (red, in the center) until the tether breaks and then these move as free exosomes (red, black outline). The larger green and blue circles depict the motion of the bulk of particles and only a few individual particles are shown. b) The output as number of particles remaining within the circle (*d* = 0.76 µm) over which the intensity was measured for all data shown. The loss of particles in time under the scenario where all particles move as untethered exosomes at a rate of 6.5 µm^2^/s (green), all particles move as CD63 does in the plasma membrane at a rate of 0.04 µm^2^/s (blue), or all are tethered with a tether that has a half-life of 8.0 seconds (red). The data is shown as grey circles and the photobleaching rate as grey squares. c) The mean of each the experimental data for temperature shown with the best fit from simulations (n = 5). Simulation parameters for the best fits are shown in Table 1. d) The fraction of each component was varied to fit individual traces at 37°C. To recreate fusion data, individual events ranged from 0-92% tethered.

**Figure 7:**
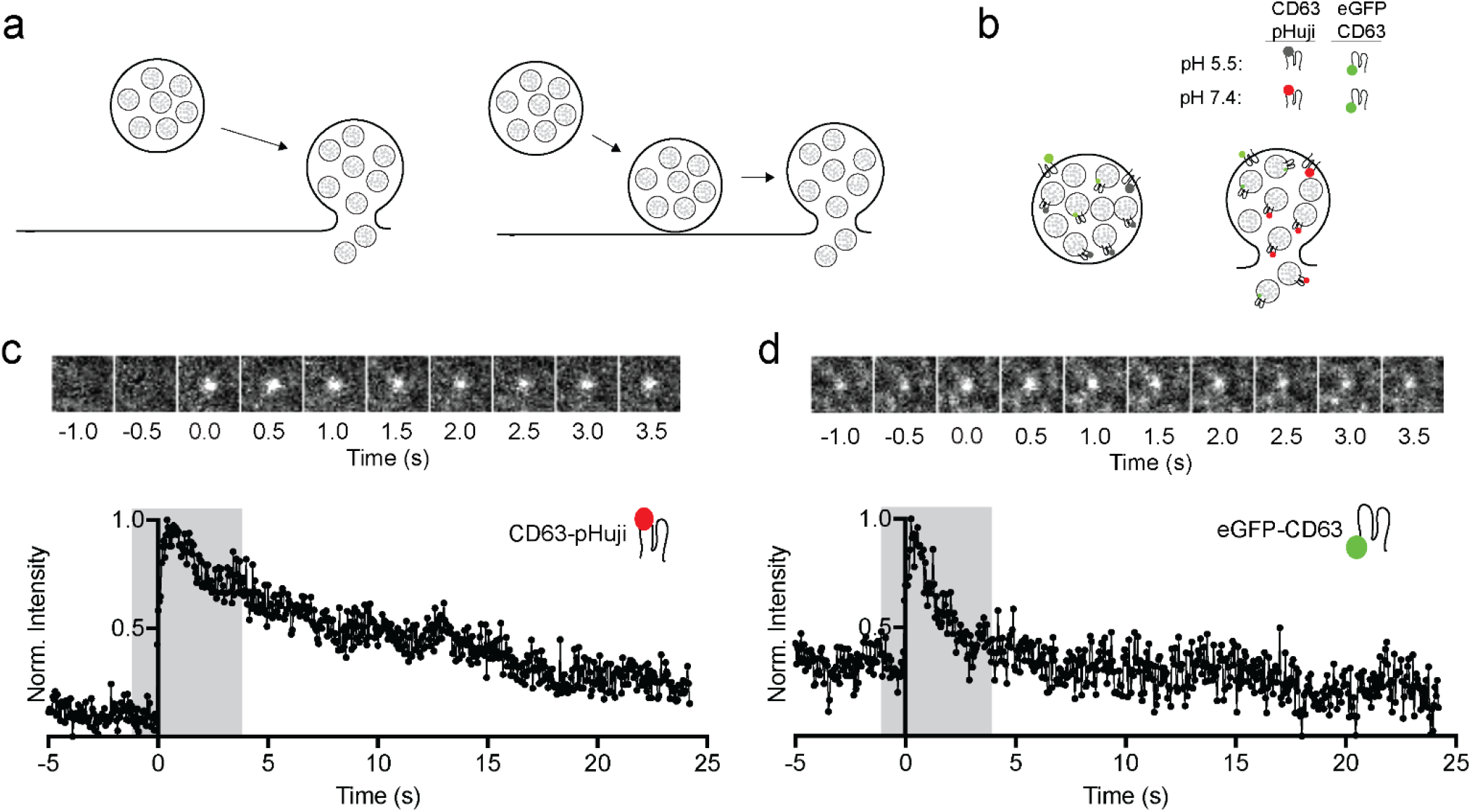
MVEs dock prior to fusion. a) Diagram illustration of crash fusion (left), and docking and fusion (right), b) Depiction of assay used to detect MVE docking prior to fusion. In crash fusion, the eGFP-CD63 vesicle marker is not present prior to fusion, however, if vesicles dock eGFP-CD63 is present before fusion. c) Single images and quantified intensity plots for a fusion event in the CD63-pHuji channel, d) Single images and quantified intensity plots of the same region during fusion of the eGFP-CD63 channel.

At all temperatures, the loss of fluorescence from the fusion site could be modeled with these three components. When comparing the simulation results to the average data at all temperatures, the variables in the simulation were: t_1/2tether_, the fraction of free exosomes, and the fraction of CD63 on the endosomal membrane, which was constrained to be 0.3+/−0.1. The fraction of attached exosomes was set to be the remaining amount of CD63 that is not on the endosomal membrane or free exosomes. The best fit for the average data at each temperature is summarized in Table 1 and shown in Fig 6C and residuals are shown in Supplemental Fig S8. The t_1/2tether_ trended longer with lower temperatures and fraction of attached exosomes also increased with lower temperatures. The amount of CD63 on the endosomal membrane was relatively constant for all temperatures and aligned well with past work that showed an endosomal fraction of 0.3-0.34 ^43^.

**Table 1:** Parameters for simulated fusion events best fits. Fusion events were simulated to best match the average intensity loss trace at each temperature. T_1/2tether_, the fraction attached, and the fraction free were unconstrained. The fraction on the endosomal membrane was constrained to 0.3+/−0.1. See Methods for details.

| | $D(\mu\text{m}^2/\text{s})$ | 23°C | 27°C | 32°C | 37°C |
| --- | --- | --- | --- | --- | --- |
| $T_{1/2\text{tether}} (s)$ | -- | 40 | 10 | 8.5 | 8 |
| *Fraction attached exosomes* | 0 | 0.6 | 0.5 | 0.5 | 0.4 |
| *Fraction free exosomes* | 6.5 | 0.08 | 0.15 | 0.15 | 0.24 |
| *Fraction endosomal* | 0.4 | 0.32 | 0.35 | 0.35 | 0.36 |
| *N events* | -- | 86 | 83 | 77 | 98 |

The individual fusion event traces at 37°C were also simulated to characterize the heterogeneity in MVE fusion that would be missed by looking at the average alone. All traces were modeled, both the biphasic and single-phase types. The t_1/2tether_ was set to 8s, as determined from the average, however, the fraction of CD63 on the endosomal membrane was varied as there is likely a distribution of CD63 localization from MVE to MVE. The fraction of CD63 on the endosomal membrane ranged from 0 to 0.6 (Fig 6D, blue) and the fraction free and tethered exosomes (Fig 6D green and red, respectively) also varied widely to accommodate the distribution of kinetics observed from single traces (examples shown in Supplementary Fig S3). Overall, the kinetics of individual MVE fusion events are highly variable but can be modeled with three types of motion for CD63-pHluorin loss from the fusion site.

### MVEs are docked prior to fusion

The steps prior to membrane fusion were also probed to determine if stable docking occurs. Whether fusion happens with newly arrived MVEs (Fig 7a, left) or MVEs that are docked before fusion (Fig 7A, right) was investigated as it can potentially point towards different mechanisms of fusion regulation. While CD63-pHuji is only visible upon the onset of fusion and serves as a fusion marker, EGFP-CD63 can be observed prior to fusion^22,27^. Using MVEs marked with both CD63-pHuji and EGFP-CD63, MVEs were visible before and during fusion (Fig 7B). In the CD63-pHuji channel (Fig 7C), fluorescence is not visible before fusion but these events were used to locate fusion sites. In the GFP-CD63 channel, green fluorescence is observed prior to fusion. Visual assessment of vesicle docking revealed that 87 of 94 events showed a visible vesicle docked at the site of fusion from the start of the movie which is at least 1 second, but generally longer. MVEs are typically docked prior to the membrane prior to fusion.

## DISCUSSION

In this work we describe an automated analysis for locating MVE fusion events from TIRF microscopy time series data and use it to characterize single MVE fusion events in A549 cells. To temporally characterize the fusion events, images were taken at a high frame rate (10-20 Hz) as events spontaneously occurred. Fusion events were located by a sharp, transient, increase in fluorescence that occurs upon a somewhat bright background since the CD63-pHluorin probe is not entirely contained within MVEs, but also appears on the cell membrane. To reduce this background, difference movies were calculated before plotting a maximum projection to locate fusion events (Fig 1, S1 and S2). This highlighted fusion events as well as other transient events where non-acidified, fluorescent vesicles moved near the cell surface or docked (Fig 2). The intensity trace of the events in time, measured from the raw data, allowed for differentiation between the types of events detected. Fusion was marked by a rapid increase in fluorescence and an exponential decay, however, diffusion and docking events were slower to rise, decayed differently or had a faster diffusion coefficient when tracked (Fig 2). Although this work focused on MVE fusion, the docked and moving events could be separately assessed in the future. This approach is capable of high throughput, automated detection and characterization of membrane fusion events and adds to currently available protocols for fusion analysis^42,44^.

The fusing MVEs ranged in size from 200-800 nm in A549 cells. Others have reported that MVE size depends on the cells and organism. In HeLa cells, EM data show that MVE range from 400 to 600 nm in diameter^27^. Using super-resolution fluorescence methods, MVEs were measured to be 1 µm in diameter in MDA-MD-231 cancer cells ^16^ and MVEs are 400 nm on average in *C. elegans* epithelial cells, based on EM analyses^45^. In the work here, MVEs secreted from A549 cells were on average 440 nm (Fig 2). It should be noted, however, that the size is calculated at the first observed moment of fusion when the MVE membrane has fused with the plasma membrane. One limitation of this approach is that expansion and loss of fluorescent content could occur during the 50 ms exposure time; this would lead to overestimation of MVE size but our measurements largely agree with EM measurements^27^, suggesting that not much fluorescence expansion occurs during the exposure time.

Prior to fusion, most MVEs dock at the plasma membrane (Fig 7). In this work, the EGFP-CD63 labeled vesicles were visible prior to fusion and only vesicles observed to undergo fusion, using a red fluorescent pH sensitive probe (CD63-pHuji), were measured. Out of 94 observed fusion events, 87 were visibly docked prior and this largely agrees with past work where CHO cells and murine embryonic primary fibroblasts were treated in ways to increase the intracellular Ca^2+^ levels to stimulate fusion^22^.

After docking, MVEs can fuse with the plasma membrane to release content and the release of content depends on temperature. Although A549 cells undergo MVE fusion at any temperature, higher temperatures increase the frequency of fusion events (Fig 5F) and the rate at which CD63 is lost from the fusion site (Fig 5A). In one respect, this is similar to stimulated fusion of synaptic vesicles, where temperature increases the number of stimulated fusion events in a variety of cell types^46,47^. However, temperature has little effect on the rate which soluble content is released and primarily affects fusion pore dilation, which occurs on a millisecond time scale in synaptic vesicle fusion^46^. The cargo released in stimulated fusion is primarily soluble, small molecules and peptides (*i*.*e*. neurotransmitters) and tethering is not expected. In MVE fusion, the time exosomes remain attached to the cell membrane accounts for the main differences observed at different temperatures (Table 1, Fig 5E and 6C); the initial loss of fluorescence, although significantly different between all temperatures, is small (Fig 5D).

Two approaches were taken to develop a mechanistic understanding of the post-fusion kinetics; experimental data was both fit and simulated using a diffusion-based model. During MVE fusion, CD63-pHluorin is likely released in at least two different ways: as exosomes and by diffusion from the endosomal limiting membrane into the plasma membrane. The working model of how fluorescence leaves the fusion site is depicted in Fig 8. Based on both the fitting and the simulations of the experimental data, we conclude that the fast (< 2s) component of the decay is a combination of free exosomes leaving and CD63 diffusing from the endosomal to the plasma membrane. The rationale is as follows:

**Figure 8:**
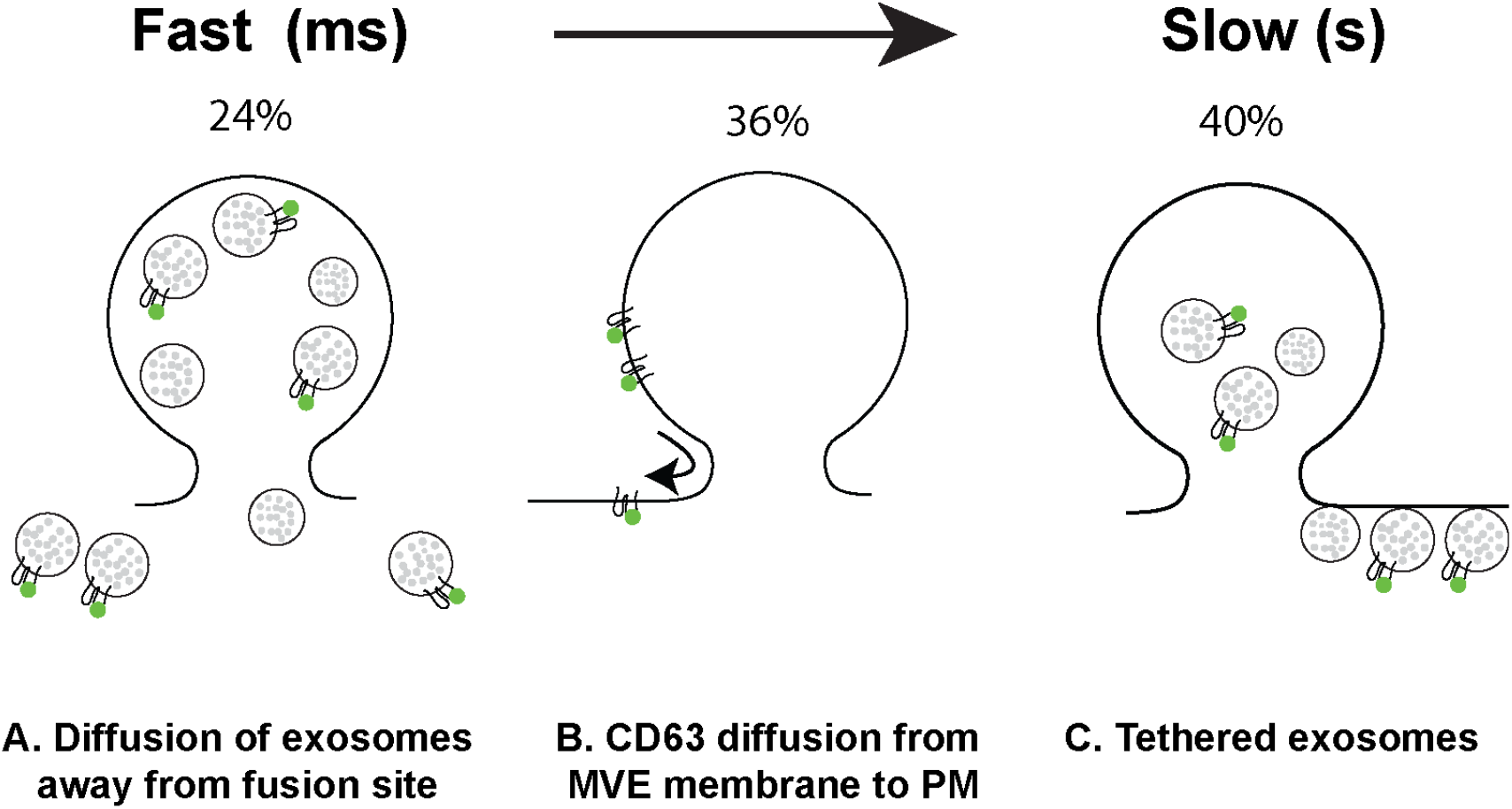
Model of how CD63-pHluorin leaves the site of fusion. A) Exosomes containing CD63-pHluorin rapidly diffuse away, < 1 s. B) CD63-pHluorin on the endosomal membrane diffuses into the plasma membrane, 1-2 s, c) Tethered exosomes containing CD63-pHluorin remain attached to the membrane and do not diffuse much. The percent of each component from a simulation of the average fusion decay is shown for 37°C.

1. Most MVE fusion events in A549 cells proceed via a biphasic decay (Fig 4). At 37°C, two thirds of the traces show biphasic decays (Fig 4D) with the faster rate on the same order of magnitude as the rate of CD63-pHluorin diffusion in the plasma membrane (Fig 4E). It is worth noting that we assume that the rate of diffusion of CD63 into the plasma membrane from the endosomal membrane is similar to the rate of diffusion of CD63 on the plasma membrane, as measured by FRAP. However, the rate of CD63 loss due to the fast component of fusion is approximately 3 times faster than expected based on CD63 plasma membrane diffusion (Fig 4e, red and blue). Here, the time it takes to leave the circle where fusion occurs, t_1/2_, is 0.40 s for the fast component and the expected time if CD63 moved at a rate of 0.039 µm^2^/s on the membrane would be 1.33 s. Other transmembrane proteins have also been noted to diffuse faster during membrane fusion. In a study of constitutive fusion using TIRF microscopy, the transmembrane protein VSVG diffused from fusion sites at a rate of 0.11 to 0.13 µm^2^/s^41,48^, whereas VSVG on the plasma membrane diffused at a rate of 0.038 µm^2^/s^49^. The diffusion of VSVG from a fusion site is 3 times faster than diffusion of the protein already present on the plasma membrane. Therefore, it is possible that the assessment of CD63-pHluorin motion on the plasma membrane using FRAP is slower than how newly delivered protein moves and this could account for the difference in the calculated rate at which protein leaves the fusion site (Fig 4E).
2. The fraction of the intensity that is lost at the faster rate (Fig 4F) also supports that this portion of the kinetics is primarily from CD63-pHluorin diffusing from the endosomal membrane into the plasma membrane. The fraction of the fast component is 0.42+/−0.02 and the fraction of CD63 reported to be on the endosomal limiting membrane by EM data is 0.30-0.34^43^. The fraction fast and likely endosomal diffusion was also constant (0.40-0.46) and not significantly different over all temperatures measured (Fig S6). In diffusion-based modeling of CD63 loss from the fusion site to simulate the decay curves, as opposed to fitting, approximately 35% of the CD63 needed to be on the endosomal membrane to recreate the experimental data (Table 1).
3. Fusion site diffusion simulations could not recreate the fast loss with *only* membrane diffusing CD63-pHluorin (Fig 6C, blue). Therefore, a small portion of CD63 needed to reside on free exosomes (8-24%, Table 1) to model the experimental data at all temperatures. The contribution of free exosomes to the fast portion could also be why the rate of release measured from fitting is faster than expected from membrane diffusion (Fig 4E, blue and red). However, fitting data with more than two exponentials was not an approach we wanted to pursue and the isolation of these two kinetic components would be challenging with the noise in our experiments. Combined, this model and the data support that CD63 leaves fusion sites by diffusing into the plasma membrane and free exosomes contribute a small amount to the initial loss

To better understand the mechanism behind the slow (> 10 s) component of the fluorescence loss post-fusion, several hypotheses were tested. First, the slow rate is almost 30 times slower than the fast at 37°C, but it is not as slow as the photobleaching rate (Fig 6C and Fig S7A-B), suggesting that CD63-pHluorin molecules are leaving the fusion site in a delayed fashion. Second, in past work on exosome secretion, others have also noted long-lived fluorescence in single fusion events and suggested this was due to tethering of exosomes^27^, although cells lacking one of the known tethers, tetherin^30^, did not remove the long-lived fluorescence^27^. Many molecules could potentially tether exosomes to the surface, such as tetraspanins, cell or matrix adhesion proteins, integrins, fibronectins and other molecules^50–52^ and attachment of exosomes to the cell surface has been noted in EM data for many years^28,29^. Exosomes themselves can act as attachment sites for migratory cells^25^, demonstrating that they are capable of sticking to the cell surface and possibly interact through the extracellular matrix. To better understand the delayed loss of fluorescence, exosome secretion was simulated as having attachments between the externalized exosome and the cell surface without knowing the identity of the attachment. To model the experimental data measured, exosomes could not remain attached eternally, however (Fig S7C-D), but instead, tethers needed to dissociate or break over the course of many seconds (8-40s). Increased temperature decreased the time for the tether to break (Table 1). Without considering the details of what molecules are at work (*i*.*e* tetherin, adhesion proteins, *etc*), this approach was able to model the long-time data well at all temperatures (Fig 6D). A model that has exosomes tethered but break in time, describes and fits the slow fluorescence decay in the experimental data well. Future experimental work in this field would help elucidate the molecules responsible for exosome attachment.

Despite our transiently tethered exosome model that fits the data well for all temperatures (Fig 6), several other mechanisms could be responsible for the observed slow fluorescence intensity loss. In neuroendocrine exocytosis, slow fusion has been measures for a number of reasons: 1) The opening and closing of a fusion pore, commonly known as “kiss and run” ^23^, significantly hinders content release. In dense core vesicle (DCV) fusion, small molecules are released first and larger peptide secretion occurs only after full fusion^53^. It is currently not known if MVEs could undergo this type of fusion, however, larger vesicles, such as MVEs, typically create stable fusion pores that dilate^54^. It is also unlikely that large molecules, like CD63-pHluorin, or exosomes would escape without the formation of a large pore. 2) The incomplete flattening of the MVE post-fusion could also give rise to slow secretion. In DCV fusion, incomplete flattening has been observed followed by endocytosis and reacidification on the order of 8 seconds in PC12 cells^55^, but much longer in chromaffin and MIN6 cells^56^. This is a viable alternative to transient tethering based on the timing observed in some neuroendocrine cells, however, more experiments are needed to characterize the shape of the MVE membrane post-fusion and the rate of endocytosis and reacidification. 3) The content of DCVs has been noted to alter the release kinetics^57,58^. Luminal proteins can affect fusion pore dilation and cause a plateau in fluorescence, like we observe in MVE fusion. 4) Finally, exosomes could temporarily stick to the glass surface used for imaging, however, different types of exosomes, for example CD9+ instead of CD63+, display fast fusion kinetics when the two were compared side by side^27^. Although a transient tether model describes the data well, permanently tethered exosomes that are quickly endocytosed should also be examined in the future.

In summary, the kinetics of MVE fusion can relate the mechanism by which fluorescently labeled exosomes leave the fusion site and the release depends on temperature. In this work, the automation of the analysis and a diffusion-based model have been developed to aid in the measurement of fusion kinetics. We conclude by proposing a model where exosomes are tethered in A549 cells; the tethers can dissociate in time and do so more readily under higher temperatures. Future studies of the role of kiss and run fusion, membrane shape changes during fusion, post-fusion endocytosis of tethered exosomes, and exosome tethering in MVE membrane fusion will further expand our understanding of how exosomes are fully released from cells.

## Author Contributions

AM collected and analyzed data and contributed to writing, AWW collected and analyzed data, ZO developed and used the simulation model, MDP developed image analyses, MTN and BLB contributed to calculations, MKK contributed to data analysis, writing, and model development.

## Acknowledgements

This work is funded by the National Science Foundation (Grants #1807455, 2122289). We thank Dinah Loerke for helpful discussions about diffusion.

## REFERENCES

1. Bebelman, M.P., Smit, M.J., Pegtel, M.D. & Baglio, R.S. Biogenesis and function of extracellular vesicles in cancer. Pharmacology & therapeutics 188, 1–11 (2018).

2. Kowal, J., Tkach, M. & Théry, C. Biogenesis and secretion of exosomes. Current Opinion in Cell Biology 29, 116–125 (2014).

3. Colombo, M., Raposo, G. & Théry, C. Biogenesis, Secretion, and Intercellular Interactions of Exosomes and Other Extracellular Vesicles. Annual Review of Cell and Developmental Biology 30, 255–289 (2014).

4. Hessvik, N.P. & Llorente, A. Current knowledge on exosome biogenesis and release. Cellular and Molecular Life Sciences 75, 193–208 (2017).

5. Thery, C., Zitvogel, L. & Amigorena, S. Exosomes: composition, biogenesis and function. Nat Rev Immunol 2, 569–79 (2002).

6. van Niel, G., D’Angelo, G. & Raposo, G. Shedding light on the cell biology of extracellular vesicles. Nat Rev Mol Cell Biol 19, 213–228 (2018).

7. Kahlert, C. & Kalluri, R. Exosomes in tumor microenvironment influence cancer progression and metastasis. J Mol Med (Berl) 91, 431–7 (2013).

8. Steinbichler, T.B., Dudás, J., Riechelmann, H. & Skvortsova, II. The role of exosomes in cancer metastasis. Semin Cancer Biol 44, 170–181 (2017).

9. Hyenne, V. et al. Studying the Fate of Tumor Extracellular Vesicles at High Spatiotemporal Resolution Using the Zebrafish Embryo. Dev Cell 48, 554–572.e7 (2019).

10. Takahashi, A. et al. Exosomes maintain cellular homeostasis by excreting harmful DNA from cells. Nat Commun 8, 15287 (2017).

11. Colvin, R.A. et al. Synaptotagmin-mediated vesicle fusion regulates cell migration. Nat Immunol 11, 495–502 (2010).

12. Proux-Gillardeaux, V., Raposo, G., Irinopoulou, T. & Galli, T. Expression of the Longin domain of TI-VAMP impairs lysosomal secretion and epithelial cell migration. Biol Cell 99, 261–71 (2007).

13. Meldolesi, J. Exosomes and Ectosomes in Intercellular Communication. Current biology : CB 28, R435–R444 (2018).

14. Vanni, I., Alama, A., Grossi, F., Bello, M. & Coco, S. Exosomes: a new horizon in lung cancer. Drug Discovery Today 22, 927–936 (2017).

15. Shen, J. et al. Advances of exosome in the development of ovarian cancer and its diagnostic and therapeutic prospect. OncoTargets and Therapy 11, 2831–2841 (2018).

16. Messenger, S.W., Woo, S.S., Sun, Z. & Martin, T.F.J. A Ca2+-stimulated exosome release pathway in cancer cells is regulated by Munc13-4. The Journal of Cell Biology 217, 2877–2890 (2018).

17. Chowdhury, R. et al. Cancer exosomes trigger mesenchymal stem cell differentiation into pro-angiogenic and pro-invasive myofibroblasts. Oncotarget 6, 715–731 (2014).

18. Clancy, J. & D’Souza-Schorey, C. Extracellular Vesicles in Cancer: Purpose and Promise. The Cancer Journal 24, 65–69 (2018).

19. Stuendl, A. et al. Induction of alpha-synuclein aggregate formation by CSF exosomes from patients with Parkinson’s disease and dementia with Lewy bodies. Brain 139, 481–94 (2016).

20. Russo, I., Bubacco, L. & Greggio, E. Exosomes-associated neurodegeneration and progression of Parkinson’s disease. Am J Neurodegener Dis 1, 217–25 (2012).

21. Rajendran, L. et al. Alzheimer’s disease beta-amyloid peptides are released in association with exosomes. Proc Natl Acad Sci U S A 103, 11172–7 (2006).

22. Jaiswal, J.K., Andrews, N.W. & Simon, S.M. Membrane proximal lysosomes are the major vesicles responsible for calcium-dependent exocytosis in nonsecretory cells. The Journal of Cell Biology 159, 625–635 (2002).

23. Stevens, C.F. & Williams, J.H. “Kiss and run” exocytosis at hippocampal synapses. Proc Natl Acad Sci U S A 97, 12828–33 (2000).

24. Shin, W. et al. Vesicle Shrinking and Enlargement Play Opposing Roles in the Release of Exocytotic Contents. Cell Rep 30, 421–431.e7 (2020).

25. Sung, B.H., Ketova, T., Hoshino, D., Zijlstra, A. & Weaver, A.M. Directional cell movement through tissues is controlled by exosome secretion. Nat Commun 6, 7164 (2015).

26. Sung, B.H. et al. A live cell reporter of exosome secretion and uptake reveals pathfinding behavior of migrating cells. Nat Commun 11, 2092 (2020).

27. Verweij, F.J. et al. Quantifying exosome secretion from single cells reveals a modulatory role for GPCR signaling. The Journal of Cell Biology 217, 1129–1142 (2018).

28. Raposo, G. et al. B lymphocytes secrete antigen-presenting vesicles. J Exp Med 183, 1161–72 (1996).

29. Simons, M. & Raposo, G. Exosomes--vesicular carriers for intercellular communication. Curr Opin Cell Biol 21, 575–81 (2009).

30. Edgar, J.R., Manna, P.T., Nishimura, S., Banting, G. & Robinson, M.S. Tetherin is an exosomal tether. Elife 5(2016).

31. Rashed, M.H. et al. Exosomal miR-940 maintains SRC-mediated oncogenic activity in cancer cells: a possible role for exosomal disposal of tumor suppressor miRNAs. Oncotarget 5, 20145–20164 (2014).

32. Hoshino, D. et al. Exosome Secretion Is Enhanced by Invadopodia and Drives Invasive Behavior. Cell Reports 5, 1159–1168 (2013).

33. Clancy, J. & D’Souza-Schorey, C. Extracellular Vesicles in Cancer. The Cancer Journal 24, 65–69 (2018).

34. Edelstein, A., Amodaj, N., Hoover, K., Vale, R. & Stuurman, N. Computer Control of Microscopes Using µManager. Current Protocols in Molecular Biology 92, 14.20.1–14.20.17 (2010).

35. Crocker, J.C. & Grier, D.G. Methods of Digital Video Microscopy for Colloidal Studies. J. Colloid Interface Sci. 179, 298 (1996).

36. Allersma, M.W., Wang, L., Axelrod, D. & Holz, R.W. Visualization of regulated exocytosis with a granule-membrane probe using total internal reflection microscopy. Molecular biology of the cell 15, 4658–4668 (2004).

37. Alnaas, A.A., Moon, C.L., Alton, M., Reed, S.M. & Knowles, M.K. Conformational Changes in C-Reactive Protein Affect Binding to Curved Membranes in a Lipid Bilayer Model of the Apoptotic Cell Surface. J Phys Chem B 121, 2631–2639 (2017).

38. Black, J.C., Cheney, P.P., Campbell, T. & Knowles, M.K. Membrane Curvature Based Lipid Sorting Using a Nanoparticle Patterned Substrate. Soft Matter 10, 2016–2023 (2014).

39. Darnton, N., Turner, L., Breuer, K. & Berg, H.C. Moving fluid with bacterial carpets. Biophys J 86, 1863–70 (2004).

40. Miesenböck, G., De Angelis, D.A. & Rothman, J.E. Visualizing secretion and synaptic transmission with pH-sensitive green fluorescent proteins. Nature 394, 192–5 (1998).

41. Toomre, D., Steyer, J.A., Keller, P., Almers, W. & Simons, K. Fusion of constitutive membrane traffic with the cell surface observed by evanescent wave microscopy. The Journal of cell biology 149, 33–40 (2000).

42. Urbina, F.L., Gomez, S.M. & Gupton, S.L. Spatiotemporal organization of exocytosis emerges during neuronal shape change. The Journal of Cell Biology, jcb.201709064 (2018).

43. Perrin, P. et al. Retrofusion of intralumenal MVB membranes parallels viral infection and coexists with exosome release. Curr Biol (2021).

44. Bebelman, M.P. et al. Real-time imaging of multivesicular body-plasma membrane fusion to quantify exosome release from single cells. Nat Protoc 15, 102–121 (2020).

45. Wehman, A.M., Poggioli, C., Schweinsberg, P., Grant, B.D. & Nance, J. The P4-ATPase TAT-5 inhibits the budding of extracellular vesicles in C. elegans embryos. Curr Biol 21, 1951–9 (2011).

46. Zhang, Z. & Jackson, M.B. Temperature dependence of fusion kinetics and fusion pores in Ca2+-triggered exocytosis from PC12 cells. J Gen Physiol 131, 117–24 (2008).

47. Oberhauser, A.F., Monck, J.R. & Fernandez, J.M. Events leading to the opening and closing of the exocytotic fusion pore have markedly different temperature dependencies. Kinetic analysis of single fusion events in patch-clamped mouse mast cells. Biophys J 61, 800–9 (1992).

48. Schmoranzer, J., Goulian, M., Axelrod, D. & Simon, S.M. Imaging constitutive exocytosis with total internal reflection fluorescence microscopy. J Cell Biol 149, 23–32 (2000).

49. Zhang, F. et al. Lateral diffusion of membrane-spanning and glycosylphosphatidylinositol-linked proteins: toward establishing rules governing the lateral mobility of membrane proteins. J Cell Biol 115, 75–84 (1991).

50. Kalra, H. et al. Vesiclepedia: a compendium for extracellular vesicles with continuous community annotation. PLoS Biol 10, e1001450 (2012).

51. Kim, D.K. et al. EVpedia: an integrated database of high-throughput data for systemic analyses of extracellular vesicles. J Extracell Vesicles 2(2013).

52. Keerthikumar, S. et al. ExoCarta: A Web-Based Compendium of Exosomal Cargo. J Mol Biol 428, 688–692 (2016).

53. Barg, S. et al. Delay between fusion pore opening and peptide release from large dense-core vesicles in neuroendocrine cells. Neuron 33, 287–99 (2002).

54. Zhang, Z. & Jackson, M.B. Membrane bending energy and fusion pore kinetics in Ca(2+)-triggered exocytosis. Biophys J 98, 2524–34 (2010).

55. Taraska, J.W., Perrais, D., Ohara-Imaizumi, M., Nagamatsu, S. & Almers, W. Secretory granules are recaptured largely intact after stimulated exocytosis in cultured endocrine cells. Proc Natl Acad Sci U S A 100, 2070–5 (2003).

56. Ohara-Imaizumi, M. et al. Monitoring of exocytosis and endocytosis of insulin secretory granules in the pancreatic beta-cell line MIN6 using pH-sensitive green fluorescent protein (pHluorin) and confocal laser microscopy. Biochem J 363, 73–80 (2002).

57. Weiss, A.N., Anantharam, A., Bittner, M.A., Axelrod, D. & Holz, R.W. Lumenal protein within secretory granules affects fusion pore expansion. Biophys J 107, 26–33 (2014).

58. Abbineni, P.S., Bittner, M.A., Axelrod, D. & Holz, R.W. Chromogranin A, the major lumenal protein in chromaffin granules, controls fusion pore expansion. J Gen Physiol 151, 118–130 (2019).

