## Supplementary Figures for "Exosome secretion kinetics are controlled by temperature"

The supplementary information contains the following figures:


| Section | Topic | Page |
|---|---|---|
| Supplementary Methods | Exosome collection and slot-blot | 2 |
| Supplementary Figure 1 | Schematic of Fusion Event Analysis | 3-4 |
| Supplementary Figure 2 | Effects of difference images and filtering | 5 |
| Supplementary Figure 3 | Examples of MVE Fusion Events | 6 |
| Supplementary Figure 4 | Slot blot of CD63 in collected exosomes | 7 |
| Supplementary Figure 5 | FRAP of CD63-pHluorin | 8 |
| Supplementary Figure 6 | Fraction Fast at all temperatures | 8 |
| Supplementary Figure 7 | Photobleaching and Simulations with fixed tethers | 9 |
| Supplementary Figure 8 | Residuals from average traces and simulations | 10 |

**Supplementary Methods:**

EV Collection and Slot-Blot Methods:

A549 cells were seeded 24 hours before exosome collection in cell culture flasks (CellTreat, Life Science Products, Frederick, CO, USA) and allowed to reach 70 – 80% confluency. After 24 hours, cells were washed 3x in DPBS containing calcium and magnesium (ThermoFisher Scientific, Item # 14040141), and incubated in DMEM (phenol red free, ThermoFisher Scientific, Item # 21063029) for 60 minutes. After the incubation time, adhered cells were scraped and collected in 1000 ul of DPBS for the total cell blot and exosome-containing supernatant was collected, and centrifuged at 300 *g* for 5 minutes, 1000 *g* for 10 minutes and 10,000 *g* for 10 minutes (2x), following the protocol described in Figure 1 of Messenger *et al* [1]. The supernatant was filtered onto nitrocellulose membrane using slot blot membrane (Bio-Rad Laboratories) and run according to the manufacturer's protocol. 5 ul (for every 50 ul of sample) of 1XBE1 buffer (40 mM HEPES, 300 mM NaCl, pH 7.5) was added to each sample before filtering onto the slot blot. CD63 was probed with the primary antibody anti-CD63 (Santa Cruz Biotechnologies, sc-5275, 1:1000) and a secondary anti-mouse antibody (Santa Cruz Biotechnologies, sc-516178, 1:1000). Then imaged on xx imager.

All other methods for supplementary figures are described in the main text.

**Supplementary Figures:**

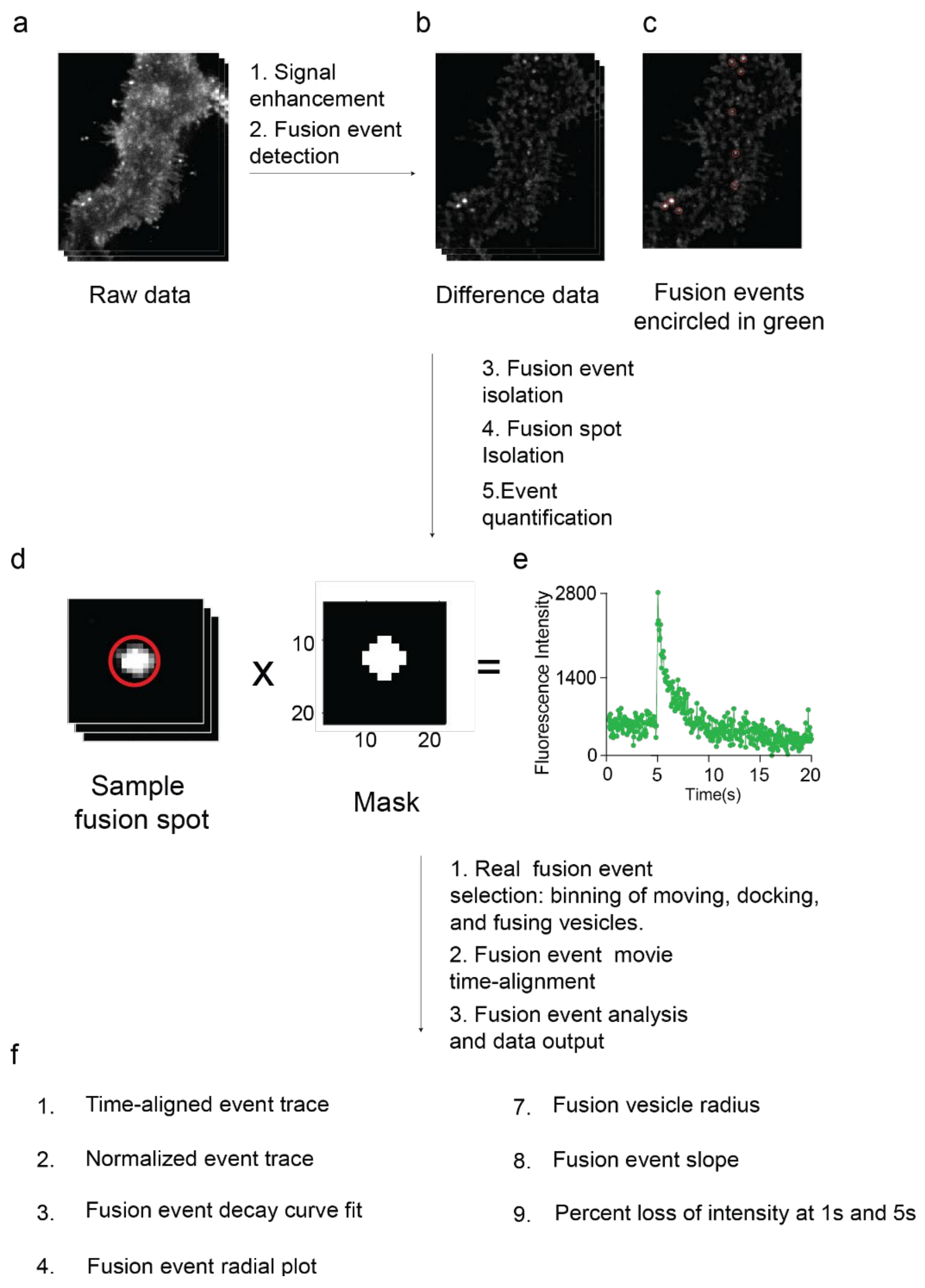


**Figure S1: Schematic overview of the automated data processing and analysis used**. A) Sample image of raw movie data. Raw data is processed, and the signal-to-noise ratio of fusion events is increased by calculating difference movies so fusion events appear as bright spots on the cell. B) Sample image of a difference movie (of raw data shown in a), showing decrease in background noise and enhanced bright spots representing fusion events. C) Maximum projection of difference movie (shown in B). From the difference movies, a maximum projection is calculated, fusion events are detected by locating the x and y coordinates of bright spots on the cell and are isolated by cropping 2.7 μm x 2.7 μm individual fusion regions. D) Sample cropped fusion event, with event circled in green (left), a mask centered on fusion event (right), and quantified fluorescence intensity plot for the fusion event over the duration of the movie (right). The mask, centered on the fusion event, is first multiplied with the potential fusion event to output intensity signal for the fusion event, excluding surrounding background signal. And the calculated output fluorescence intensity is then averaged, to obtain the average time course intensity of the fusion event. Real fusion event peaks are filtered from non-real events (see Methods and Figure S3 for more details), and both fusion event movies and traces are aligned by artificially setting the peak at t = 0 s. F) List of various analyses output by the algorithm. Following time alignment, fusion events are analyzed, and a variety of results related to the kinetics, vesicle size and the number of events per minute are output. (Cell movie# 1100AM)

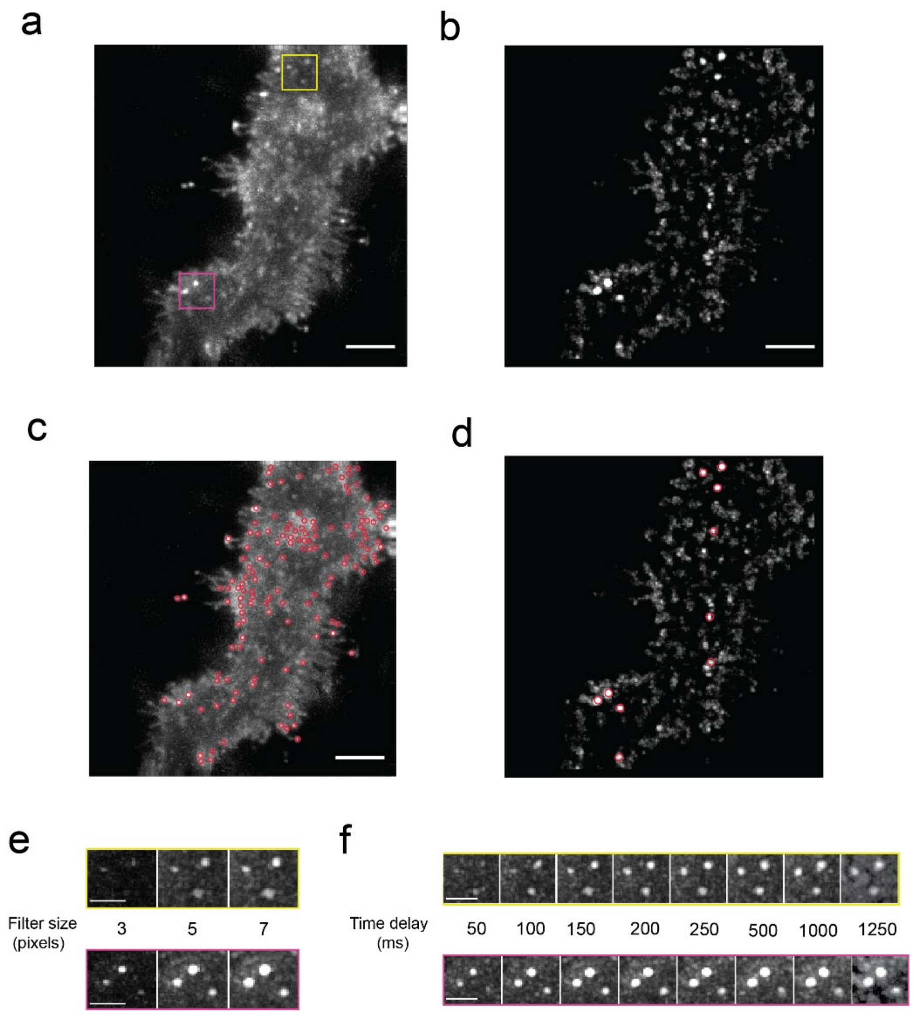


**Figure S2: Differential movies are calculated to enhance the fusion events relative to stationary vesicles** (Steps 1-2 in Figure S1). A) A single image of a transiently transfected A549 cell expressing CD63-pHuji with 50 ms exposure. Six fusion event spots are highlighted by pink and yellow boxes. B) A maximum projection of a difference movie was generated using a time interval of 25 frames (1.25 s) and bright spots indicate changes in signal, possibly fusion events. There is a decrease in the plasma membrane background after calculating the difference movie. C) Approximately 140 bright spots were detected using a maximum projection of the raw data, locations are overlaid on the maximum projection of the raw movie. D) Ten bright spots were detected using the difference image shown in B. The regions are overlaid on the maximum projection of the difference movie. Note that the third feature from the top was not detected in C. Scale bar in A-D is 4.36 µm and 2 µm in E and F. E) Maximum projection images of difference data showing changes in fusion event S/N (for events highlighted in A) after applying a bandpass filter of 3, 5, or 7 pixels. The difference movie was calculated using time interval of 150 ms and then bandpass filtered with the sizes shown in pixels, where one pixel is 0.109 µm. F) Maximum projection images showing changes in fusion event S/N (for events highlighted in A) upon application of a bandpass filter of 7 pixels and different time delays. (Cell movie # 1100AM)

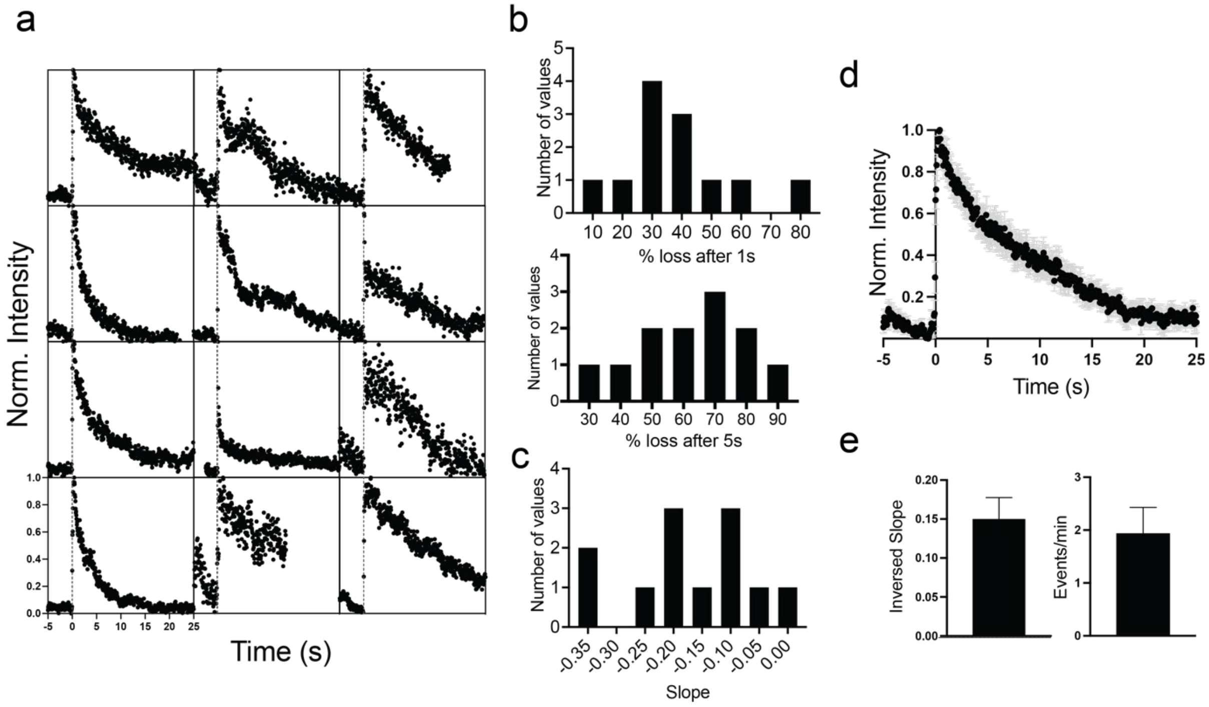


**Figure S3: Example of individual fusion events from two cells.** a) Example of 12 normalized fusion events from two cells where time lapse movies were taken repeatedly on the same cells. Two events were cut short (top row, right and bottom row, middle) and could not be fit well with an exponential function. b) Histogram showing distribution of percent lost after one second (top) and five seconds (bottom) post fusion. c) Histogram showing slope distributions of data in a. d) Average fusion event trace of the 12 events in a. e) Average Inversed slope (left) and average events per minute per cell (right). The error bar is the SEM variation from movie to movie. (Cell movies #1496AM-1507AM, 1952AM-1956AM)

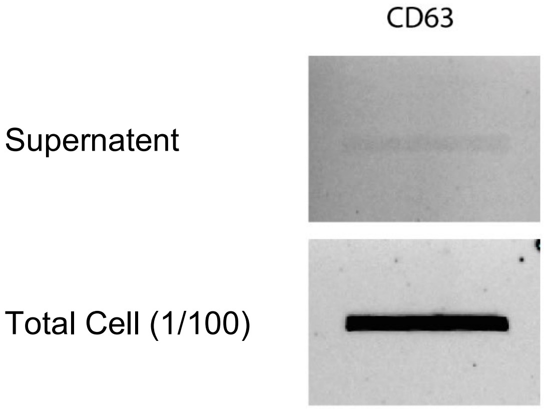


**Figure S4: CD63 is present on A549 EVs.** A549 cells were cultured to 70-80% confluency then the cell media was replaced with DMEM (no phenol red) and incubated over a 30 minute period. The media was collected and centrifuged as described in the supplemental methods. For the total cell fraction, cells were scraped from the dish in DPBS. Both the total cell fraction and EV containing supernatant were run on a slot blot and probed for CD63. Slot blot results were confirmed by at least three independent replicates.

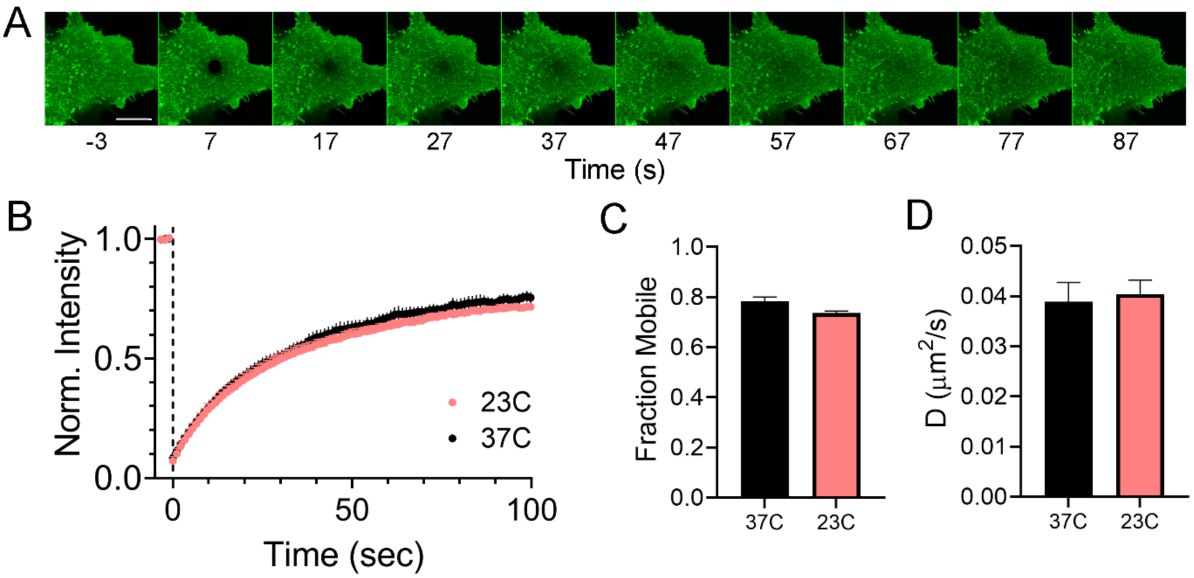


**Figure S5: Diffusion rate of CD63-pHluorin in plasma membrane are similar at 23C and 37C.** A) Montage showing photobleaching and recovery of CD63-pHluorin on plasma membrane of A549 cells. Confocal image of the bottom of the cell before photobleaching (-3s) is followed by the recovery process at intervals of 10s. Scale bar = 10 µm. B) Intensity of the bleached region in time at 23°C and 37°C (n=10 cells each). C) The mobile fraction at 23°C and 37°C. D) Diffusion coefficient for CD63 at 23°C and 37°C. Standard error is shown in all figures. There are no significant differences in plots C and D.

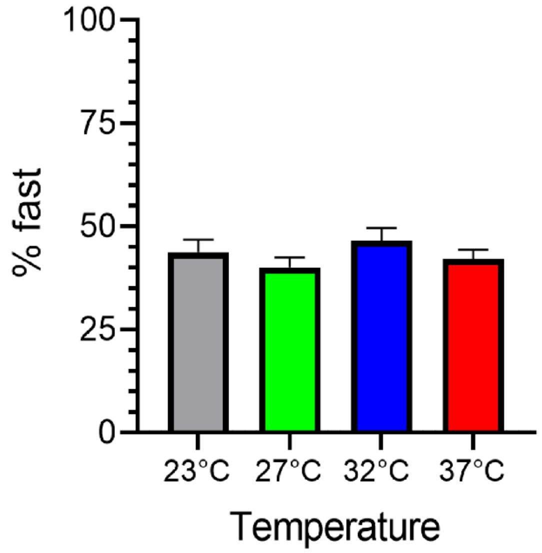


**Figure S6: Temperature dependence of the fraction fast in two phase fusion events.** Temperature does not change the distribution between fast and slow components in fusion event decays that contain two components. None are statistically different, and the fraction fast varies from 40-46%

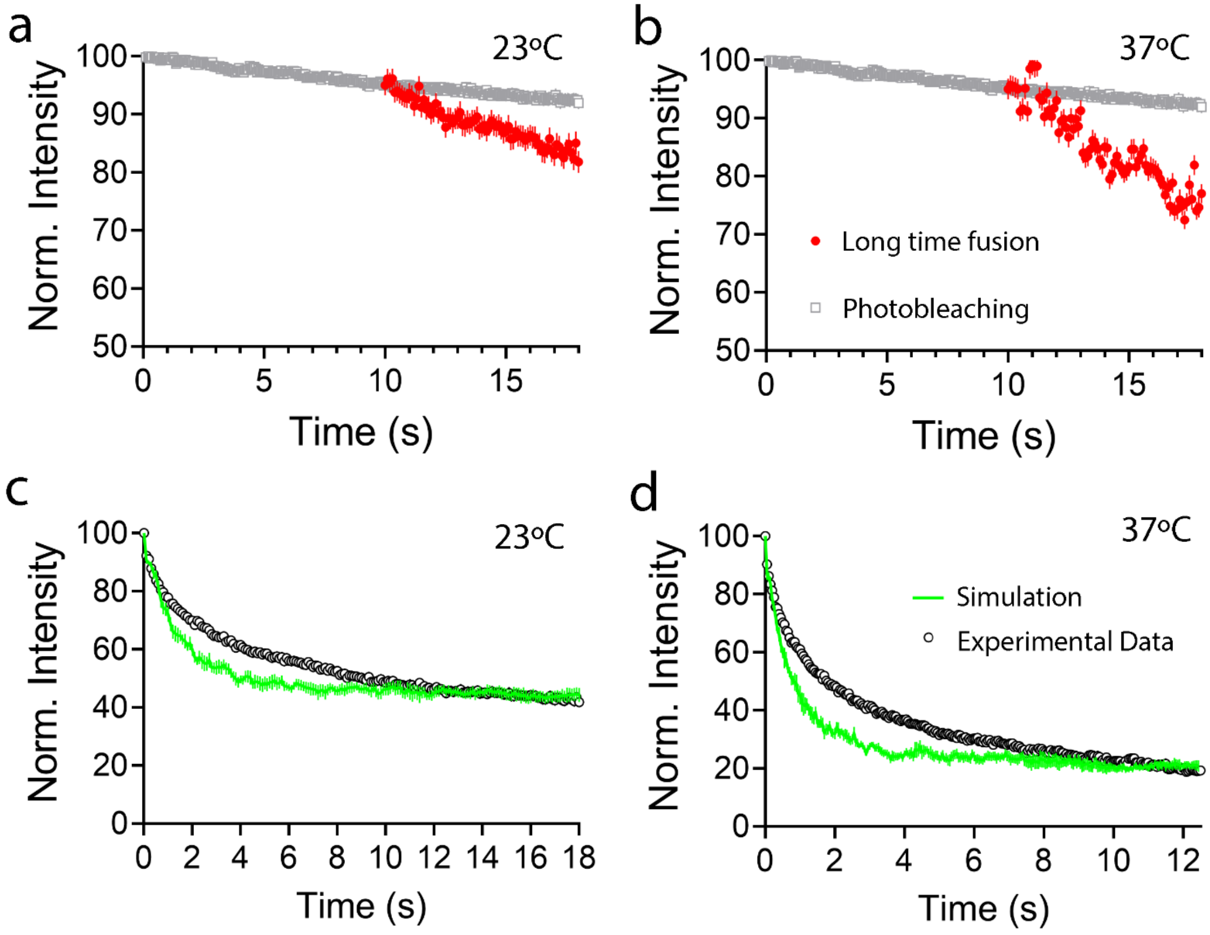


**Figure S7: Experimental data cannot be fit well to the rate of photobleaching or unbreakable tethers.** The long-time fluorescence intensity loss decays faster than the photobleaching rate at A) 23°C and B) 37°C. The experimental data in A and B were renormalized to align to the photobleaching intensity at 10s for ease of comparison. Comparison of the average, experimental fusion event decay curve and a simulation of fusion data in presence of unbreakable tethers at C) 23°C and D) 37°C. The simulation was chosen where the initial decay and the plateau were best aligned based on residuals.

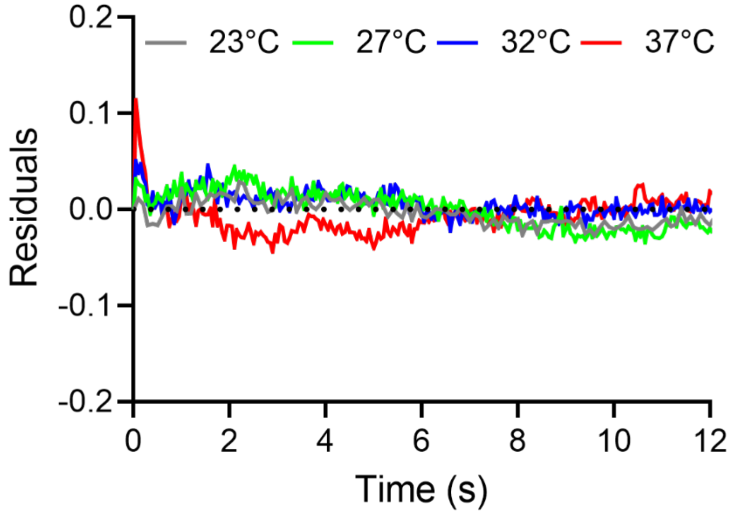


**Figure S8: The residuals from the simulation for modeling all temperatures when compared to the experimental data.** The residuals were calculated by subtracting an average of five or more simulations from the average intensity of single fusion events for each temperature. Each temperature was fit by varying the time of the tethers to break, the fraction of free exosomes, and the fraction of endosomal CD63. The remaining fraction of CD63 that was not free or endosomal was considered tethered.

**Supplementary References:**

[1] Messenger SW, Woo SS, Sun Z, Martin TFJ. A $Ca^{2+}$-stimulated exosome release pathway in cancer cells is regulated by Munc13-4. J Cell Biol. 2018 Aug 6;217(8):2877-2890. doi: 10.1083/jcb.201710132. Epub 2018 Jun 21. Erratum in: J Cell Biol. 2019 Apr 1;218(4):1423. PMID: 29930202; PMCID: PMC6080937.
